# Experience Alters the Timing Rules Governing Synaptic Plasticity and Learning

**DOI:** 10.1101/2022.11.28.518128

**Authors:** Sriram Jayabal, Brandon J. Bhasin, Maxwell Kounga, Jennifer DiSanto, Aparna Suvrathan, Mark S. Goldman, Jennifer L. Raymond

## Abstract

The brain learns about the statistical relationships between events in the world through associative synaptic plasticity, controlled by the timing between neural events. Here, we show that experience can dramatically alter the timing rules governing associative plasticity and learning. In normally reared mice, the timing requirements for short- and long-term associative plasticity at synapses in the oculomotor cerebellum are precisely matched to the 120 ms delay for visual feedback to the circuit about behavioral errors. This specialization of the plasticity rules for the constraints of a particular circuit and learning task is acquired through experience. In dark-reared mice that never experienced visual feedback about oculomotor errors, synapses defaulted to a coincidence-based plasticity rule, with a corresponding delay in the timing of learned eye movements. This temporal metaplasticity persists into adulthood; when mice reared normally from birth were moved to dark housing as adults, the task-specific timing requirements for plasticity and the temporal accuracy of learning were lost and then re-established when visual experience was restored. Computational modeling suggests two general classes of biologically plausible mechanisms, each with multiple possible implementations, that can tune plasticity to distinct features of the statistics of neural activity. Temporal metaplasticity provides a potentially general mechanism for experience-dependent improvement in the way a circuit solves the “temporal credit assignment problem” inherent in most learning tasks, thereby providing a candidate synaptic mechanism for meta-learning.

## INTRODUCTION

The timing between neural events controls the induction of plasticity at most synapses, providing a mechanism for learning about the correlations between events in the world, for example, between actions and their consequences. There is considerable variation across synapses in the intervals between neural events that are effective for inducing plasticity, ranging from a few tens of milliseconds to as long as ∼1 s.^1–14^ In a few cases, a precise correspondence has been found between the timing contingencies for plasticity at certain synapses and specific biophysical, circuit-level or behavioral constraints on learning, raising the possibility that variations in the timing rules for plasticity are not random, but rather reflect functional specializations for particular learning tasks.^3,5,7,11,12^ Here we tested the role of experience in generating functional specialization of plasticity rules.

One of the most striking specializations of the timing rules for plasticity identified to date is at the cerebellar parallel fiber-Purkinje cell synapses.^3^ During learning, parallel fiber-Purkinje cell synapses that cause errors undergo associative synaptic depression, triggered by feedback about errors signaled by the single climbing fiber input to each Purkinje cell.^15^ The feedback delay for climbing fibers to signal errors varies across functional regions of the cerebellum, from a few tens of milliseconds for climbing fibers carrying somatosensory feedback^16^ to greater than 100 ms for visual feedback,^3,17^ and potentially longer for climbing fibers signaling cognitive errors.^18^ For optimal learning, the timing contingencies for plasticity would need to compensate for the relevant feedback delay for climbing fibers to signal errors, so that synapses that were active at the time an error was generated are selectively weakened. Consistent with this possibility, the most effective pairing interval between parallel fiber and climbing fiber activity for inducing associative synaptic depression varies considerably for parallel fiber synapses onto different Purkinje cells.^3^ Moreover, in the cerebellar flocculus, which supports oculomotor learning, a precise match was found between the timing requirements for synaptic plasticity and the feedback delay in the circuit. When an eye movement fails to perform its function of stabilizing images on the retina, the delay between the parallel fiber activity generating the errant eye movement and the visual climbing fiber activity signaling the oculomotor error, as reflected by image motion on the retina, is approximately 120 ms *in vivo.*^3,17,19,20^ In slices of the cerebellar flocculus, both long- and short-term depression is selectively induced by pairing parallel fiber activation with climbing fiber activation at this functionally relevant delay of 120 ms, and not by pairing intervals of 100 or 150 ms or coincident pairing.^3^ Here, we report a major role of experience in generating this specialization of the timing rules for plasticity for the functional requirements of the circuit and learning task.

## RESULTS

### Experience alters the timing rules for synaptic plasticity

To assess the role of experience in generating plasticity rules matched to the feedback delay in the oculomotor circuit, mice were dark-reared to eliminate experience of the normal, 120 ms delay for visual feedback about oculomotor errors to be reported to the oculomotor cerebellum (cerebellar flocculus) by the climbing fibers. The timing rules for plasticity at the parallel fiber (PF)-Purkinje cell synapses were then measured in slices of the cerebellar flocculus of the dark-reared mice, and compared with those in slices from control mice reared on a normal 12 h dark/12 h light schedule. If the plasticity rules in the cerebellum are shaped by experience, then the normal or abnormal inputs to the cerebellum from other brain areas, reflecting the normal or dark-rearing experience, should yield different plasticity rules in the cerebellum of mice reared in these different conditions.

Previous studies have shown that the induction of LTD at the PF-Purkinje cell synapses is influenced by the parallel fiber-climbing fiber (PF-CF) pairing interval along with a number of additional experimental parameters,^21^ including the number of PF stimuli used in each PF-CF pairing,^22^ the number of CF stimuli used in each pairing,^23,24^ the number of pairings,^25^ the presence of inhibition,^26,27^ and extracellular factors such as calcium concentration^23^ each of which could vary dynamically *in vivo*.^28^ Here, to isolate plasticity at the excitatory PF-Purkinje cell synapses, inhibition was blocked. To characterize the timing requirements for LTD induction, single PF and CF stimuli were delivered at a fixed pairing interval in each experiment, since the use of trains of PF or CF stimuli would create multiple PF-CF timings, complicating the analysis of timing-dependence. To facilitate comparison of *in vitro* and *in vivo* results, PF-elicited spiking in the Purkinje cells was used to assess plasticity, in addition to PF-elicited synaptic responses in the Purkinje cells. To ensure that any difference in the rules for plasticity reflects the *in vivo* rearing experience of the animals, all experimental parameters were identical in slices from normally reared and dark-reared mice.

The results revealed a major effect of experience on the timing requirements for associative plasticity. In slices of the cerebellar flocculus of normally reared mice, LTD of PF-elicited spiking in the Purkinje cells was selectively induced by a PF-CF pairing interval of 120 ms, and not by coincident pairing (**Fig. 1A***, red,* **Fig. S1**), as previously reported for PF-elicited EPSCs and PF-elicited EPSPs in Purkinje cells.^3^ In mice dark-reared from birth, the timing requirements for LTD were altogether different than in normally reared mice. In dark-reared mice, the functionally relevant, 120 ms PF-CF pairing interval induced no LTD of parallel fiber elicited spiking (**Fig. 1B***, red,* **Fig. S1, S2**) indicating that the ability of delayed climbing fiber feedback to induce LTD is only acquired through experience. In the absence of normal visual experience, LTD seems to default to a coincidence-based rule. In the dark-reared mice, coincident activation of parallel fibers and climbing fibers induced LTD (**Fig. 1B**, *blue*), whereas in normally reared mice coincident pairing was completely ineffective at inducing LTD (**Fig. 1A**, *blue*). Similarly, LTD of PF-elicited synaptic currents (EPSCs) in the Purkinje cells was selectively induced by coincident PF-CF pairing (**Fig. 1C**, *blue*), and not by a 120-ms pairing interval in slices from dark-reared mice **(Fig. 1C**, *red*), contrary to what is observed in normally reared mice.^3^

**Figure 1.**
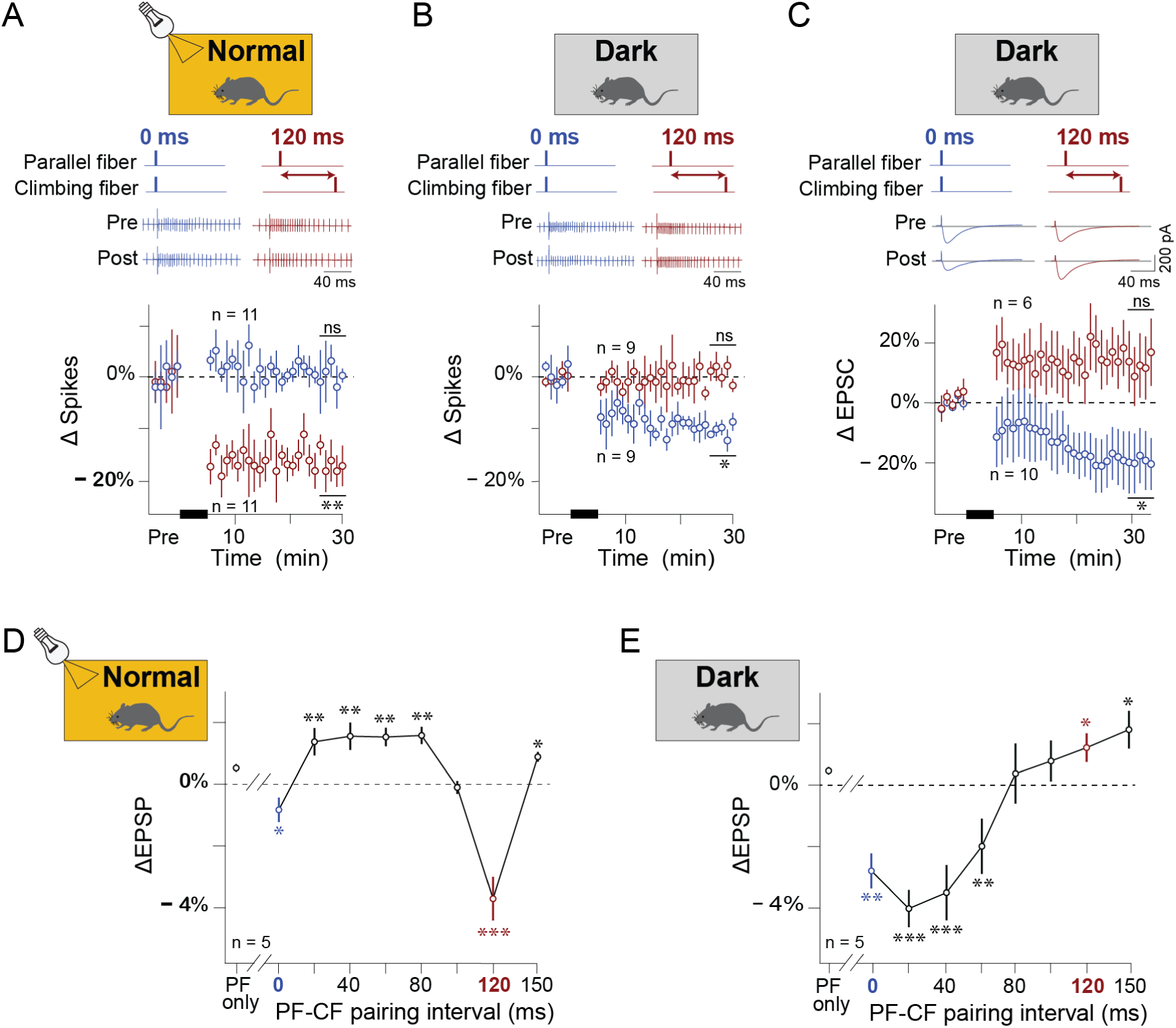
Experience alters the timing rules for associative plasticity at the parallel fiber-Purkinje cell synapses. (**A,B**) Long-term depression (LTD) of parallel fiber (PF)-elicited spiking in Purkinje cells in slices of the cerebellar flocculus from mice reared normally in cycles of 12 hr light/12 hr dark (**A**) or dark-reared from birth (**B**). ***Top,*** Examples of PF-elicited spiking in a Purkinje cell before (*pre*) and after (*post*) paired stimulation of the PF and climbing fiber (CF) inputs to the Purkinje cell (300x at 1 Hz) with a pairing interval of 0 ms (coincident; *blue*) or 120 ms (*red*). ***Bottom***, Average change in PF-elicited spiking in Purkinje cells induced by PF-CF pairing, delivered at time indicated by *black bar*, normalized to pre-pairing baseline. In normally reared mice (**A**), LTD of PF-elicited spiking was induced by a PF-CF pairing interval of 120-ms (*red*; \*\**P* = 0.001, single-sample *t*-test, *n* = 11 cells) but not 0 ms (*blue*; *P* = 0.863; single-sample *t*-test, *n* = 11 cells). In dark-reared mice (**B**), LTD was induced by a PF-CF pairing interval of 0 ms (*blue*; \**P* = 0.013, single-sample *t*-test, *n* = 9 cells) but not 120 ms (*red*; *P* = 0.226, single-sample *t*-test, *n* = 9 cells). Error bars in all panels are S.E.M. (**C**) LTD of PF-elicited excitatory postsynaptic currents (EPSCs) in Purkinje cells, in dark-reared mice. ***Top***, Example EPSCs before and after PF-CF pairing. ***Bottom***, Average change in EPSC amplitude, normalized to pre-pairing baseline. LTD was induced by a PF-CF pairing interval of 0 ms (*blue*; \**P* = 0.0347, single sample t-test, n = 10 cells) but not 120 ms (*red*; *P* = 0.0781, Wilcoxon signed-rank test, n = 6 cells). (**D,E**) Short-term depression (STD) of PF-elicited excitatory postsynaptic potentials (EPSPs) in Purkinje cells induced by a single PF-CF pairing in normal (**ci**) and dark-reared (**cii**) mice. Average change in EPSP amplitude is plotted as a function of the PF-CF pairing interval. *Blue* (0 ms) and *red* (120 ms) data points correspond to the PF-CF pairing intervals used to test LTD in panels **A,B** and **C**. The rearing condition altered the timing contingencies for short-term plasticity (interaction effect, rearing x pairing interval, P < 0.0001; effect of rearing, *P* = 0.0003; effect of pairing interval, P < 0.0001; 2-factor ANOVA following align-and-rank transformation of data using ARTool(41,42)). \**P <* 0.05, \*\**P <* 0.01, \*\*\**P* < 0.001, single sample t-test for plasticity at each pairing interval.

Associative short-term depression (STD) exhibited experience-dependent changes in its timing contingencies (**Fig. 1D**) similar to those observed for LTD. In cerebellar slices from normally reared mice, a single parallel fiber-climbing fiber pairing induces associative STD of PF-Purkinje cell synapses lasting a few seconds, with timing requirements similar to those for LTD.^3,29^ The rapid decay of STD allows multiple PF-CF pairing intervals to be compared at the same PF-Purkinje cell synapses, permitting a fine-grained characterization of the dependence of associative plasticity on the PF-CF pairing interval.

Short-term plasticity was assessed by comparing PF-elicited synaptic potentials (EPSPs) in the Purkinje cells measured 1 s before and 1s after a single PF-CF pairing. As in previous work,^3,29^ PF-elicited EPSPs were used to assess short-term plasticity rather than PF-elicited EPSCs to avoid rapid switching between current-clamp for pairing and voltage-clamp for measurement, which is hard on the health of cells. In mice reared with normal visual experience, STD of the parallel fiber-Purkinje cell synapses in the flocculus was selectively induced by the functionally relevant 120 ms pairing interval, and not by pairing intervals of 100 or 150 ms (**Fig. 1D**, *red,*^3^). This timing requirement was robust to a two-fold change in extracellular calcium (**Fig. S3).** In contrast, in dark-reared mice, STD was selectively induced by shorter pairing intervals of 0 to 60 ms (**Fig. 1E**). Most notably, the 120 ms PF-CF pairing interval that corresponds to the feedback delay in the circuit was completely ineffective at inducing STD in the dark reared mice, whereas it induced robust STD in normally reared mice (compare *red* in **Fig. 1D vs E**).

Together, the results for both short- and long-term plasticity, measured with three different readouts of neural plasticity–parallel fiber-driven synaptic currents (EPSCs), synaptic potentials (EPSPs) and spiking in Purkinje cells–provide convergent evidence that the specialized timing rules governing cerebellar plasticity are not simply pre-determined, but are altered by experience. Experience suppressed the ability of coincident neural activity to drive associative plasticity, and created a new ability of delayed climbing fiber activation to induce plasticity. We refer to this novel capacity of synapses to undergo experience-driven changes in the timing rules for plasticity as temporal metaplasticity.

### Behavioral correlate of temporal metaplasticity

Specialization of the timing requirements for associative plasticity has the potential to influence the temporal accuracy of learning. The plasticity rules in normally reared mice should enable the behavioral errors reported by climbing fibers to selectively modify only those synapses that were active at the earlier time when the error was generated, solving what is widely referred to as the temporal credit assignment problem.^30,31^ In contrast, the coincidence-based plasticity rule observed in the dark-reared mice should cause climbing fiber feedback to depress synapses active around the time an error is *reported* to the flocculus, rather than depressing synapses active at the earlier time when the error was *generated*. Given that Purkinje cell firing drives eye movements with a precision of 3-5 ms in the relationship between Purkinje cell firing rate and eye velocity,^32^ temporal metaplasticity would be expected to alter the timing of learned eye movements. To test this prediction, we characterized the temporal accuracy of oculomotor learning in the normally and dark-reared mice.

Optokinetic reflex (OKR) adaptation is a form of oculomotor learning supported by LTD at the parallel fiber-Purkinje cell synapses in the cerebellar flocculus.^33,34^ With one hour of training, normally reared mice learned to increase the velocity of their eye movements to better stabilize an optokinetic visual stimulus on the retina by matching their eye velocity to that of the stimulus (**Fig. 2A-D**, *gold*). Dark-reared mice were also capable of oculomotor learning, even though the OKR training session was their first visual experience (**Fig. 2A-D**, *green*). However, there was a striking difference in the timing of the learned eye movements in the dark-reared mice compared to the normally reared mice. In dark-reared mice, the learned eye movements were delayed by approximately 100 ms relative to the visual stimulus and relative to normally reared mice (**Fig. 2C-E; Fig. S4**), a time shift comparable to the altered timing requirements for associative plasticity measured at the synaptic level (**Fig. 1**). Thus, the temporal metaplasticity driven by normal visual experience is accompanied by an improvement in the temporal accuracy of learning.

**Fig. 2.**
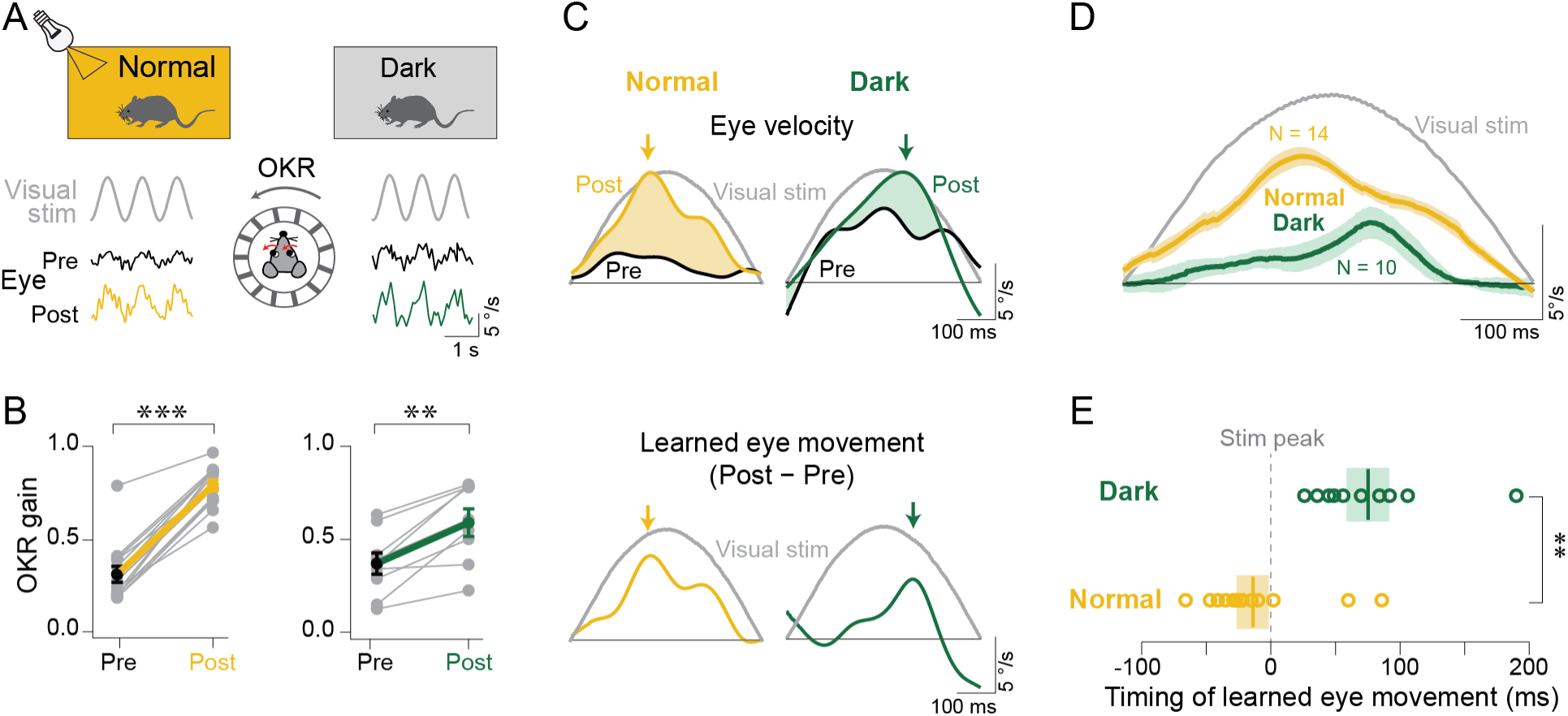
Experience improves the temporal accuracy of learning. (A) During optokinetic reflex (OKR) adaptation, mice learn to more accurately track the motion of a striped visual stimulus with their eye movements. Raw traces show example horizontal eye velocity responses to a sinusoidally moving visual stimulus (i.e., the OKR) at the beginning (*Pre*) and end (*Post*) of a 60-min training session, in representative normally reared (***left***) and dark-reared (***right***) mice. (B) One hour of OKR training induced a learned increase in the OKR gain (ratio of eye velocity to visual stimulus velocity) in both normally reared (***left; \*\*\*****P* < 0.0001, matched pairs Wilcoxon signed rank test) and dark-reared (***right*; \*\****P* = 0.0036, matched pairs *t*-test) mice. *Grey* symbols show results for individual mice, averages across mice are shown in *black* and *gold* (normal) or *black* and *green* (dark-reared). The baseline OKR gain before the training (*Pre*) was similar in normal and dark-reared mice (*P* = 0.3610, Wilcoxon test). Error bars indicate S.E.M. (C) ***Top***, Eye movement responses, averaged across 10-45 cycles of the optokinetic stimulus, in an individual normally reared mouse (***left***) and a dark-reared mouse (***right***), pre- and post-training. ***Bottom*,** The learned component of the eye movement response, isolated by subtracting the pre-training eye movement response from the post-training response (equal to the shaded *gold* and *green* areas in ***top*** panels). (D) Average learned component of the eye movement response to an optokinetic visual stimulus. Traces show the mean ± S.E.M. across normally reared mice (*gold; N* = 14) and dark-reared (*green; N* = 10) mice. (E) Timing of the learned eye movement (time of peak learned eye velocity relative to peak optokinetic stimulus velocity), calculated for each mouse (*circles*) was delayed in dark-reared mice (*green)* relative to normally reared mice (*gold*) (\*\**P* = 0.0002, independent samples *t*-test), and relative to the visual stimulus (*P* = 0.0007, single-sample *t*-test). *Solid vertical lines*: average peak learned timing across animals; *shaded region*: ± S.E.M.

### Temporal metaplasticity in adulthood

Experience could alter the timing rules for plasticity, not only during development, but also in adulthood. After normal rearing for the first 60 days of life, to allow the timing rules for cerebellar plasticity and the temporal accuracy of learning to develop normally (**Fig. 3A,B, *top***), mice were moved to dark housing conditions to eliminate further experience of visual feedback about oculomotor performance. At 20-day intervals, OKR learning was assessed, and a progressive degeneration of the temporal accuracy of learning was observed (**Fig. 3A**, *green*). After 60 days in the dark, the eye movements learned in response to OKR training were delayed by more than 100 ms (**Fig. 3A**, *dark green*), comparable to the delay in the learned eye movements of mice dark reared from birth (**Fig. 2C-E**). Extending the dark experience an extra 60 days had no additional effect on the temporal accuracy of learning (**Fig. S5**).

**Fig. 3.**
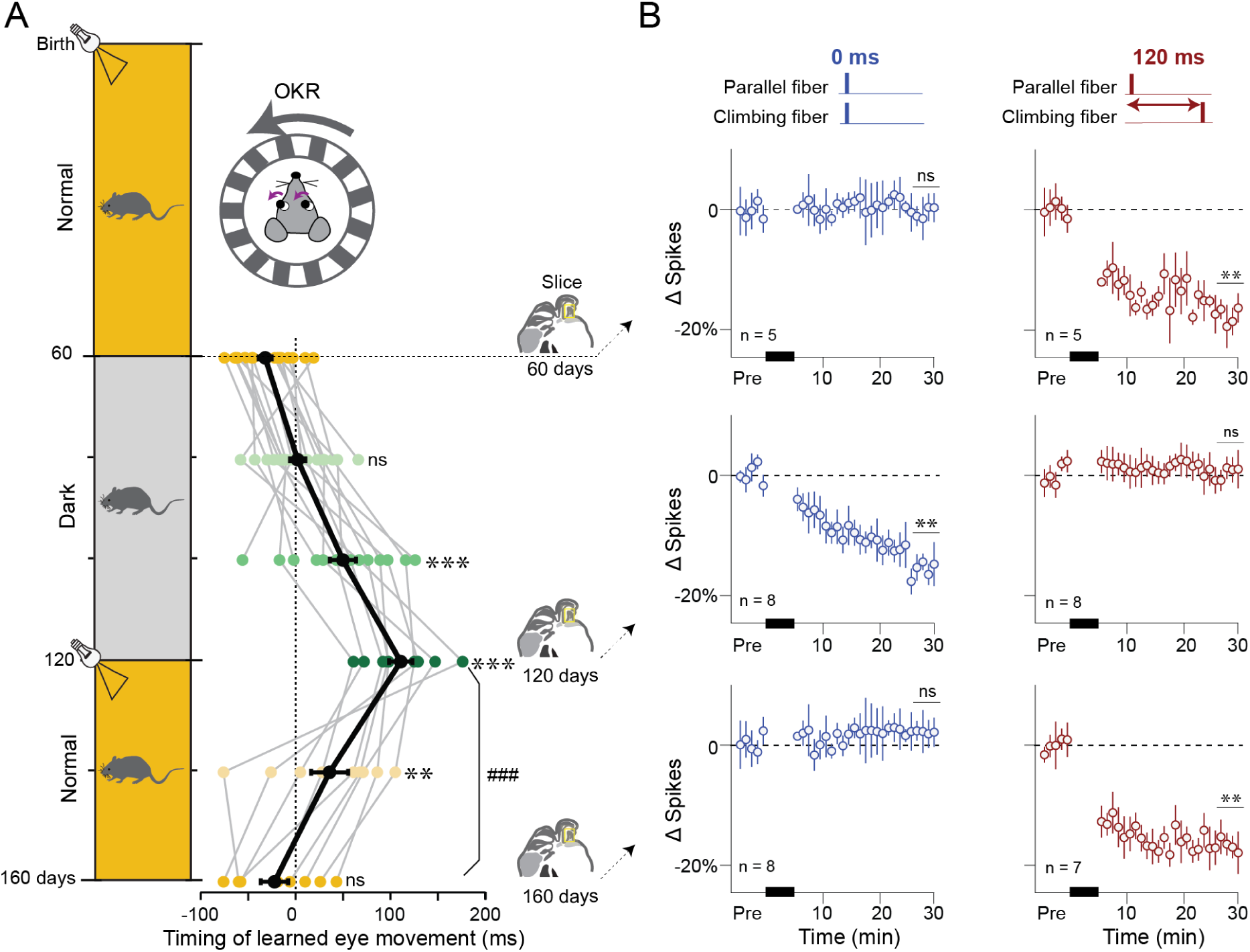
Temporal metaplasticity persists into adulthood. (A) ***Left,*** Schematic showing the timeline of manipulating visual experience during adulthood. Mice were normally reared on a 12 hr light/12 hr dark cycle for 60 days post-birth. After this period of normal development, mice were maintained in darkness for the next 60 days (postnatal days P60-P120), followed by a return to normal visual experience for an additional 40 days (P120-P160). OKR learning was tested at 20-day intervals, starting at P60. Slice electrophysiology experiments were performed at P60, P120, or P160. ***Right,*** Timing of the learned eye movement responses (time of peak learned eye velocity relative to peak velocity of the optokinetic stimulus) in individual mice tested at 20 day intervals (*grey lines* connecting *colored symbols*) and the average across mice (*black*). Dark experience during adulthood induced a progressive delay in the timing of the learned eye movements (F_5,74_ = 18.24, *P* = <0.0001, ANOVA; P60 vs P100 and P60 vs P120, \*\*\**P* < 0.0001, Tukey). Following return to normal visual experience, the timing of the learned eye movements was restored to baseline (P60 vs P140, \*\**P* < 0.002; P60 vs P160, *P* = 0.99; P120 vs P160, ^###^*P* < 0.0001; Tukey). (B) Long-term depression (LTD) of parallel fiber (PF)-elicited spiking in Purkinje cells in slices of the cerebellar flocculus. ***Top***, After 60 days of normal rearing from birth, LTD of PF-elicited spiking was not induced by coincident pairing of PF and climbing fiber (CF) stimulation (0 ms pairing interval, 300x at 1 Hz) (*blue, P* = 0.45, single-sample *t*-test), but LTD was induced by a PF-CF pairing interval of 120 ms (*red*, \*\**P* = 0.004, single-sample *t*-test). ***Middle***, Following 60 days of dark experience during adulthood, from P60-P120, LTD tested at P120 was induced with a 0 ms PF-CF pairing interval (*blue,* \*\**P* = 0.007, single-sample *t*-test), and not a 120 ms pairing interval (*red*, *P* = 0.58, single-sample *t*-test). ***Bottom***, After return from dark to normal visual experience for 40 days (P120-P160), LTD tested at P160 was not induced with a 0 ms PF-CF pairing interval (*blue, P* = 0.32, single-sample *t*-test), but LTD was induced with a 120 ms pairing interval (*red*, \*\**P* = 0.005, single-sample *t*-test). Error bars in all panels are S.E.M. Number of cells is given in lower ***left*** of each panel. *Black bars* indicate the time of PF-CF pairing.

In parallel to the behavioral changes, the timing rules for neural plasticity underwent changes in response to altered experience in adulthood. Guided by the behavioral findings, the timing rules for plasticity were assessed in slices of the cerebellar flocculus of mice that experienced 60 days of normal rearing followed by 60 days of dark experience. In these mice, associative LTD was induced by coincident pairing of parallel fiber and climbing fiber activation, but not by the 120 ms pairing interval that is effective in normally reared mice (**Fig. 3B, *middle row***). Thus, even after the normal timing rules for plasticity are established during development (**Fig. 3B, *top row***), these rules are actively maintained by visual experience, and in the absence of that experience, the timing rules revert to the coincidence-based plasticity observed in synapses of mice dark-reared from birth (**Fig. 1**). If, after 60 days of dark experience during adulthood, mice were then returned to normal housing and visual experience, the temporal accuracy of OKR learning recovered to normal in about 40 days (**Fig. 3A, *bottom***), as did the ability of the functionally relevant 120 ms pairing interval to induce LTD (**Fig. 3B, *bottom***).

Thus, normal visual experience was not only essential for establishing the timing rules for plasticity during development, but also for maintaining and reestablishing the timing rules for plasticity in the adult. In other words, the capacity for temporal metaplasticity was retained in adulthood, with correlated changes in the temporal accuracy of learning.

### Spike-timing statistics can drive adaptive changes in the timing requirements for plasticity in a computational model of the cerebellar circuit

Our experiments establish that the timing requirements for plasticity are altered by experience and tightly aligned with the functional requirements of the specific circuit and behavioral task. In principle, a synaptic plasticity rule matched to the 120 ms feedback delay for climbing fibers to signal oculomotor errors could be genetically pre-specified. Alternatively, an adaptive process could tune the timing requirements for plasticity to become matched to the circuit’s characteristic delay. We used computational modeling to test the plausibility of the hypothesis that temporal metaplasticity is an adaptive process.

#### Two mechanistic classes for adjusting the timing rules for plasticity based on the timing of presynaptic and postsynaptic spiking

Just as the timing of presynaptic and postsynaptic spikes drives changes in synaptic weight according to a plasticity rule (**Fig. 4A**;^8,35–38)^, we propose two mechanisms by which spike timing could simultaneously drive changes in the plasticity rule itself according to a temporal metaplasticity rule.

**Figure 4.**
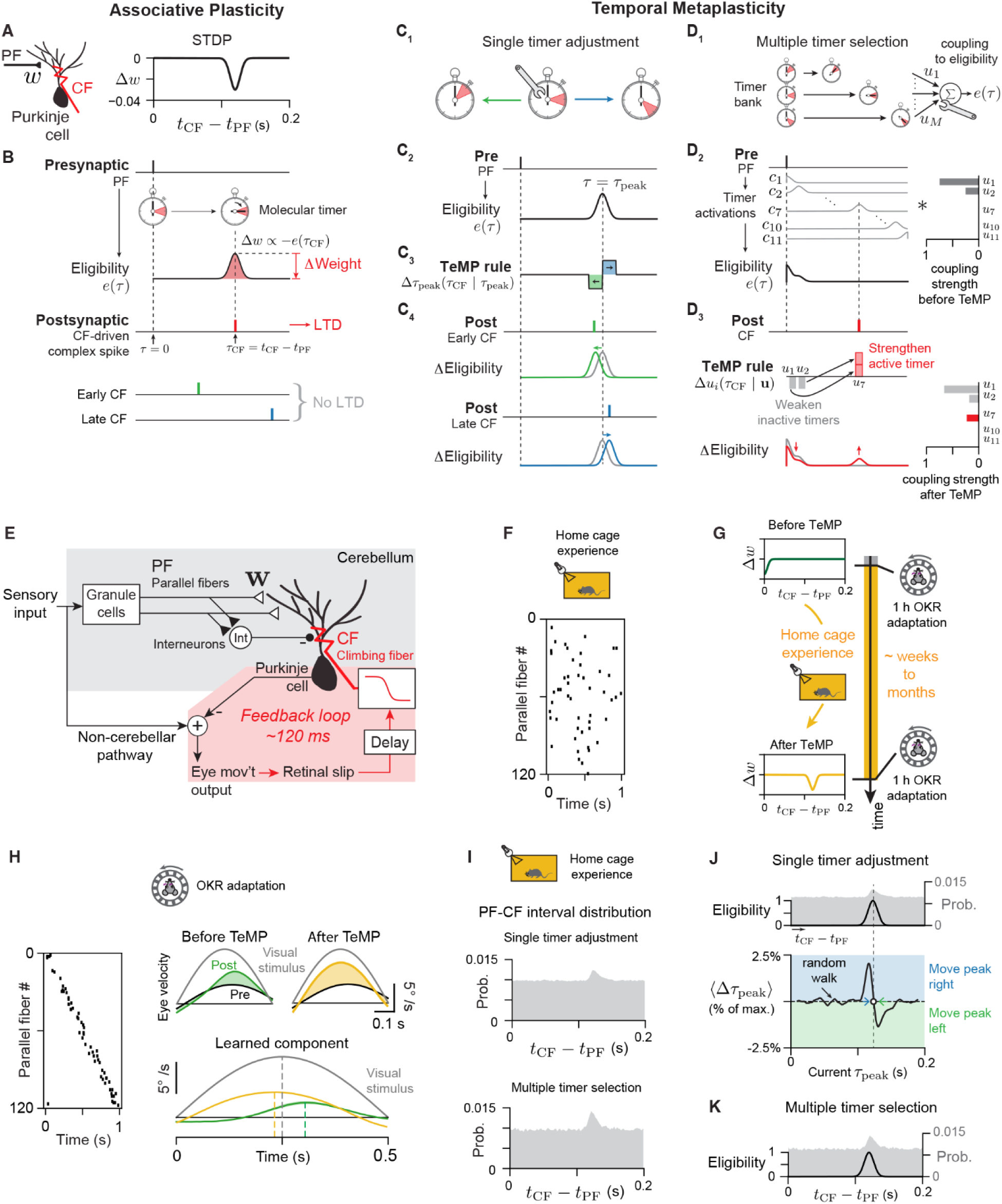
Models of candidate mechanisms for temporal metaplasticity. **(A)** Spike-timing dependent plasticity (STDP) window for associative LTD at the parallel fiber-Purkinje cell synapse. A decrease *Δw* in the synaptic weight *w* occurs when the time *t_CF_* of the CF-elicited postsynaptic complex spike in the Purkinje cell follows the time *t_PF_* of a parallel fiber (PF) input spike within a narrow eligibility window centered around 120 ms. **(B)** Schematic of spike-timing dependent associative LTD at parallel fiber-Purkinje cell synapses. A PF input spike (*Presynaptic*) at time τ = 0 starts a molecular timer (*stopwatch*) that creates a time-varying eligibility window *e*(τ) for plasticity. The change in weight *Δw* at that PF synapse is proportional to the value of the eligibility (*e*(τ); *shading in stopwatch*) at the time τ_CF_ of the CF-elicited postsynaptic complex spike in the Purkinje cell (*hand of stopwatch*). A complex spike within the eligibility window (*red*) causes LTD, whereas complex spikes outside the eligibility window (*green, blue*) cause no LTD. **(C, D)** Two possible classes of mechanisms for temporal metaplasticity: adjustment of a single timer, and selection from a bank of timers. (**C_1_, C_2_**) In the single timer adjustment mechanism (**C_1_**), the eligibility window *e*(τ) for associative plasticity is generated by a single molecular timer initiated by presynaptic (“Pre”) input (**C_2_**). (**C_3_**) The peak timing τ_peak_ of the eligibility window can be shifted earlier or later according to the value of a temporal metaplasticity (TeMP) window at the time of the postsynaptic spike (*TeMP rule*). (**C_4_**) If the CF-triggered postsynaptic spike occurs before (*green*) or after (*blue*) the peak eligibility, the time of peak eligibility is shifted slightly earlier or later, respectively. (**D_1_**) In the timer selection mechanism, eligibility for plasticity results from a weighted sum of activities of a bank of molecular timers that peak at different times after presynaptic input (**D_1_**, *left*). Temporal metaplasticity is accomplished by changing the coupling strength (contribution), *u_m_*, of each molecular timer to the eligibility for plasticity (**D_1_**, ***right***). (**D_2_**) Example eligibility trace (***bottom***) is constructed by the product of the activations *c_i_* of many timers (***left***) and the coupling strengths *u_i_* (***right***). (**D_3_**) At the time of a postsynaptic (CF) spike, the coupling strength of the timer most active at that time is strengthened, and the couplings of the other timers are decreased (*TeMP rule*). This results in an updated eligibility trace (***bottom left***, *red*) and set of coupling strengths (***bottom right***). (**E-K**) Oculomotor circuit model with integrated plasticity and metaplasticity. (**E**) Architecture of cerebellar circuit model. Eye movement output was linearly related to Purkinje cell activity. Each Purkinje cell was driven by 120 parallel fibers (PFs) and simultaneously inhibited by a molecular layer interneuron (MLI) that pools PF activity. Oculomotor errors, defined as motion of the optokinetic stimulus relative to the eye (retinal slip) probabilistically elicited climbing fiber (CF) spikes with a mean delay of 120 ms, which in turn drove both temporal metaplasticity and associative LTD of the weights, **w**, of parallel fiber-Purkinje cell synapses. (**F**) PF spiking was assumed to be uncorrelated over the prolonged duration of home cage experience (but highly similar results were found when PF spiking was strongly correlated; see **Supplementary** Fig. 6). (**G**) Timeline of simulations follows the experimental timeline. Simulations started with an untuned (coincident) plasticity rule. Following a prolonged duration of home cage experience, plasticity assessed using a 1 h OKR adaptation experiment showed that the timing rules for plasticity shifted to the 120 ms PF-CF interval characteristic of the feedback delay in the circuit. (**H**) Effect of metaplasticity on OKR learning. ***Left***, PF spiking during OKR adaptation with a 1 Hz sinusoidal stimulus. ***Top right***, Eye velocity before (*“Pre”*, *black*) and after (*“Post”*) OKR adaptation, shown for the start of the simulation (*“Before TeMP”*) and after metaplasticity has occurred (*“After TeMP”*). ***Bottom right***, the learned component of eye velocity shifts from lagging the visual stimulus, to being close to aligned with the visual stimulus, similar to the experiments (cf. **Fig. 2D, E**). (**I-K**) Temporal metaplasticity learns to adjust the eligibility trace to the peak of the PF-CF interval distribution. (**I**) PF-CF interval distributions computed from circuit model simulations with the single-timer adjustment (***top***) or multiple-timer selection rule (***bottom***) show a peak at the 120 ms circuit feedback delay on top of a uniform baseline. (**J**) Dynamics of the single-timer adjustment mechanism. ***Top***, Eligibility window (*black*) following metaplasticity aligns with the peak of the PF-CF interval distribution (*gray*). ***Bottom***, mean shift <*Δ*τ_peak_> of the eligibility window peak as a function of the current location of the eligibility window peak for the PF-CF interval distribution of panel **I**. When the peak of the eligibility window, τ_peak_, is in the flat (uniform) portion of the PF-CF distribution, it random walks. When τ_peak_ becomes close to the peak of the PF-CF interval distribution (*vertical dashed line*), it shifts (*blue and green arrows*) in a direction that aligns the eligibility window peak with the PF-CF interval distribution peak, solving the temporal credit assignment problem. (**K**) The multiple timer selection mechanism also aligns the peak of the eligibility window (*black*) to that of the PF-CF interval distribution, as described in panel **D**.

For the first class of mechanism, we conceptualize temporal metaplasticity as the adjustment of a single molecular timer that controls the timing requirements for plasticity (**Fig. 4B**, ***top***).

Presynaptic (PF) input starts the timer, which after an adjustable delay period produces a brief window of eligibility for associative plasticity (**Fig. 4B**, ***middle****; e(τ) and red region in stopwatch*). The eligibility window defines the plasticity rule: if a postsynaptic spike (for Purkinje cells, the CF-driven complex spike) occurs within the eligibility window (**Fig. 4B**, *hand of stopwatch within red region*), the synapse undergoes associative plasticity and the synaptic weight changes by an amount proportional to the magnitude of the eligibility window at the time of the spike (**Fig. 4B**, ***bottom***).

The temporal metaplasticity rule describes how the time of the postsynaptic spike changes the length of the delay period created by the molecular timer and hence changes the plasticity rule (**Fig. 4C_1_-C_3_**). If a postsynaptic spike arrives just before eligibility peaks, the delay period of the molecular timer is decreased, shifting the time of peak eligibility earlier and thus making shorter pairing intervals effective at inducing a change in synaptic weight (**Fig. 4C_3_-C_4_**, green).

Likewise, if the postsynaptic spike arrives just after peak eligibility, the time of peak eligibility is shifted later (**Fig. 4C_3_-C_4_**, blue). We refer to this class of temporal metaplasticity mechanism as the “single timer adjustment” mechanism.

In the second class of mechanism, there is a bank of molecular timers, each with a fixed time (**Fig. 4D_1_**, red shaded regions in stopwatches) at which they can contribute to eligibility for plasticity. Eligibility for plasticity is controlled by a weighted sum of the timer activations (**Fig. 4D_2_**, ***left***) with adjustable weights (**Fig. 4D_1_-D_2_**, ***right***, coupling strengths). The coupling strengths define the plasticity rule by indicating how the synaptic weight will change in response to a postsynaptic spike at the associated timer’s time of activation. The temporal metaplasticity rule then describes how the time of the postsynaptic spike changes the coupling strengths of each timer as follows: when a postsynaptic spike occurs (**Fig. 4D_3_**, **top**), the coupling of the timer that is active at the time of the spike is strengthened, and the coupling of all the other, inactive timers are weakened (**Fig. 4D_3_**, “TeMP rule”), leading to an updated set of coupling strengths and eligibility window (**Fig. 4D_3_**, ***bottom row***). We refer to this class of mechanism as the “multiple timer selection” mechanism.

#### Adjustment of the timing rules for plasticity and oculomotor behavior in a cerebellar circuit model

When implemented in a circuit model with a characteristic feedback delay, each of the two classes of metaplasticity mechanism can use the spike-timing statistics arising from normal activity to adjust the timing requirements for plasticity to match the feedback delay. We modeled a general circuit architecture reflective of circuits throughout the cerebellum receiving feedback at a delay,^3^ with specific reference to the oculomotor cerebellum for comparison with the experimental results (**Fig. 4E**; see **STAR Methods**). Neural activity in the model was generated based on signaling properties measured experimentally during oculomotor learning and, where these properties are not well known, we tested different assumptions. The sensory input that drives oculomotor behavior was represented by parallel fiber activity in the circuit model.

Purkinje cell activity was modeled as a linear combination of parallel fiber input activity. Purkinje cells in the flocculus are just two synapses from the motor neurons that move the eye, and changes in their firing rate have an approximately linear effect on the eye movement behavior (eye velocity;^32,39^). Therefore, eye movement output was modeled as a linear combination of a Purkinje cell activity plus a nonplastic contribution from a non-cerebellar pathway. Climbing fibers represented errors in the eye movement output relative to a desired eye movement and arrived at the Purkinje cells with a distribution of delays centered around the experimentally observed characteristic delay of 120 ms (^3,17^; see **STAR Methods**).

To model how temporal metaplasticity could use neural activity to match the plasticity rule to the characteristic delay of the circuit, we simulated parallel fiber activity representing the variety of sensory input that might be expected during behavior in the home cage under normal visual rearing conditions (**Fig. 4F**; see **STAR Methods** for details). Starting from an initial coincidence-based plasticity rule, spiking in the circuit could adaptively match the plasticity requirements to the characteristic delay. With a single timer, this could be accomplished by the timer adjustment rule of **Fig. 4C_3_**. With multiple timers, we used a rule in which each out-of-sync timer’s coupling is weakened by a fixed amount (if above zero), which is then transferred to the active timer (**Fig. 4D**).

To compare to the experimental paradigm (**Figs. 2, 3**), we also simulated a brief period of OKR learning with a sinusoidal visual stimulus before and after the period of home cage experience (**Fig. 4G,H**, **top**). In the model, the learned component of the OKR response prior to metaplasticity was delayed relative to that after metaplasticity by around 100 ms (**Fig. 4H**, ***bottom***). Thus, the model suggests that the differences in behavior we found experimentally (**Figs. 2, 3**) can be accounted for just by differences in the timing requirements for plasticity at parallel fiber-to-Purkinje cell synapses.

A key finding from the simulations was that temporal metaplasticity can match the plasticity rule to the characteristic delay in the circuit through experience, even though synapses experience spurious spike timings other than the characteristic delay. The distributions of parallel fiber-climbing fiber intervals experienced by each synapse over the course of the simulations consisted of a peak corresponding to the characteristic delay of the circuit on top of a uniform “background” probability arising from the chance pairing of unrelated parallel fiber and climbing fiber inputs (**Fig. 4I**). The distribution took this form both when the activity of the population of parallel fibers was uncorrelated (as in **Figs. 4F, S6A**) and when parallel fibers were strongly temporally correlated (as in **Fig. 4H**, ***left***; **Fig. S6B**).

The robust ability of the temporal metaplasticity rules to match the characteristic delay is a consequence of the rules’ dynamics. For the single-timer-adjustment rule, the average change < *Δ*τ_peak_ > in the time of peak eligibility for plasticity (**Fig. 4J**) is determined by how much the distribution of spike timings (**Fig. 4I**) overlaps the right and left lobes of the temporal metaplasticity rule (**Fig. 4C_3_**). In the simulations, the value of τ*_peak_* is initially taken to be close to zero, where the shape of the distribution of PF-CF spike intervals (**Fig. 4I**) is locally uniform. In this region, spikes are just as likely to move the peak earlier as later, causing it to diffuse (**Fig. 4J**, “random walk”). If τ*_peak_* shifts late enough to get close to the peak of the distribution, the right lobe of the metaplasticity rule will overlap more of the PF-CF interval distribution than the left lobe, leading to rightward drift of τ*_peak_* (**Fig. 4J**, blue arrow). If τ*_peak_* diffuses to a value larger than the peak of the distribution, the metaplasticity rule will similarly cause a leftward drift in τ*_peak_* (**Fig. 4J**, blue arrow). As a result, τ*_peak_* is driven toward a value near the peak of the spike-timing distribution (**Fig. 4J**, open circle).

For the multiple-timer-selection rule, the dynamics of temporal metaplasticity are defined by the changes in coupling strength for each molecular timer. The average change in coupling strength is positive for the timer that is active near the peak of the distribution, because it is most likely to be strengthened, and negative for the rest of the timers. Thus, this rule creates a winner-take-all competition among different potential timing rules for associative plasticity.

The above results may also provide insight into why the timing requirements for plasticity return to near-coincident during extended experience in the dark (**Fig. 3**). The spike-timing distribution in the dark could have a peak close to coincidence as a result of non-visual signals driving the climbing fiber^40,41^ with shorter characteristic delays. Temporal metaplasticity would then match the timing requirements for plasticity to these shorter delays during extended dark experience.

Alternatively, climbing fiber activity may become uncorrelated from parallel fiber activity in the absence of visual errors. In this case, simple modifications to either of the temporal metaplasticity rules presented above can produce a return to a default coincidence-based timing rule in the absence of a peak in the spike-timing distribution (**Fig. S7**; **STAR Methods**).

### Different biologically plausible temporal metaplasticity mechanisms extract distinct statistical features of spike-timing distributions

Many possible molecular signaling pathways could mediate temporal metaplasticity. Here, we show how temporal metaplasticity could be implemented using simple biochemical network motifs such as molecular switches, accumulators, and reversible posttranslational modifications.

#### Biochemical implementations of single timer adjustment mechanism

We implemented a molecular timer through the accumulation of a molecular species to a threshold (**Fig. 5A**, biochemical network; **Fig. 5B, dynamics of molecular activations**; **STAR Methods**;^42,43^). Presynaptic input turns on a “timer on/off” switch (Fig. 5A,B, activation of molecule A1 to active form A1*). This causes the accumulation of activated molecular species B* that, when large enough to cross a (soft) threshold, activates a molecule C* that makes the synapse eligible for plasticity (Fig. 5B). C* then turns off the “timer on/off” switch, causing the accumulated B* to degrade, and thus closing the window for eligibility.

**Figure 5.**
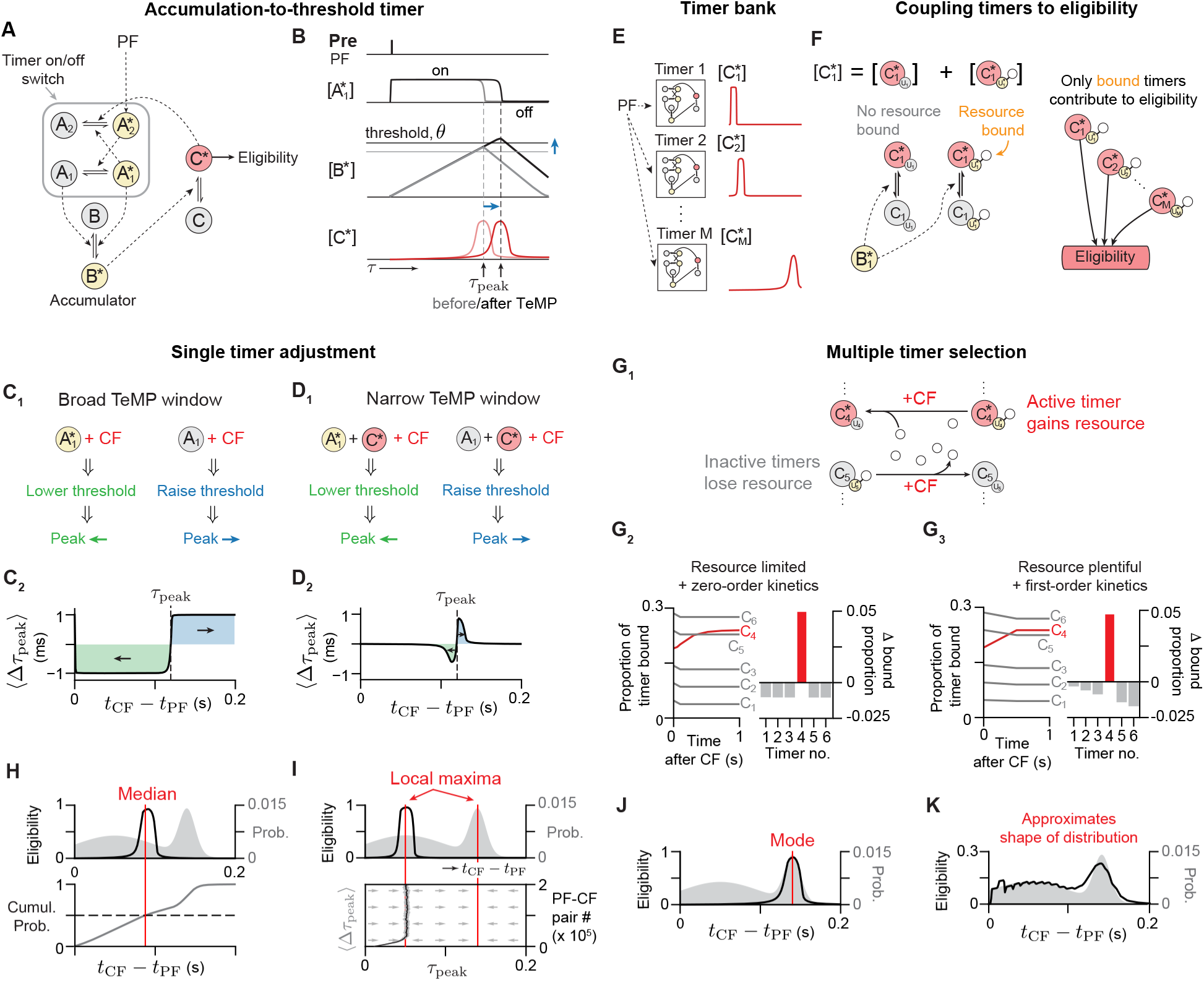
Different biochemical network implementations of the temporal metaplasticity mechanisms extract different features of the input timing interval distribution. **(A, B)** Molecular realization of a single adjustable timer, based on accumulation-to-threshold dynamics. (**A**) Reaction diagram. The timer consists of three functional components – a “timer on/off” switch created by positive feedback between the A_1_ and A_2_ molecules, an accumulator, and an eligibility molecule. Asterisks denote the active form of each molecular species. (**B**) Single timer dynamics. Parallel fiber (PF) input activates the timer on/off switch (i.e., activates A_1_*), leading to the accumulation of B*. When above an approximate threshold concentration (*horizontal lines*), B* activates the eligibility molecule C. Eligibility for plasticity (*red traces*) is controlled by the time-varying concentration of C^∗^. C* drives the inactivation of A_2_*, turning off the timer on/off switch and causing B* to be deactivated. C* then rapidly deactivates when B* falls below threshold, closing the window of eligibility. By lowering or raising the threshold for activation of C (*horizontal black and gray lines*), the time of peak eligibility τ_peak_ can be moved earlier or later in time. **(C, D)** Adjustment of the accumulation-to-threshold timer is controlled by the concentrations of A_1_, A_1_* and C* at the time of climbing fiber (CF) input. (**C_1_**) Since A_1_* is only present until the peak eligibility τ_peak_ is reached, the presence (***left***) or absence (***right***) of A_1_* at the time of the CF input indicates whether the CF input was too early (before τ_peak_) or too late (after τ_peak_), indicating that τ_peak_ should be shifted earlier (***left***, *green*) or later (***left***, *green*). (**C_2_**) Leftward or rightward shifts of the eligibility peak are accomplished by changing molecule C’s activation kinetics so as to lower or raise, respectively, its activation threshold. (**D_1_, D_2_**) To produce the version of the rule with narrow temporal metaplasticity window (**Fig. 4C_3_**), metaplasticity events are limited to the narrow/brief period of time when the eligibility molecule C* is present. **(E-G)** Implementation of the multiple timer selection mechanism. (**E**) Bank of accumulation-to-bound timers that, following a PF spike input, activate at different, fixed times determined by their concentration of molecule C_i_*. See **Supplementary** Fig. 8A for an alternative implementation. (**F**) Each eligibility molecule C_i_ has a binding domain element U_i_ that can be activated to U_i_* by binding to an eligibility-inducing resource. Each timer contributes to the eligibility at the time of its activation (i.e., when it is in form C_i_*), with amplitude given by the amount of resource it has bound (U_i_*). Diagrams simplified to capture the essence of the full binding reaction diagram of **Supplementary** Fig. 8B. (**G**) Binding and unbinding of resource is determined by the time of the CF spike. (**G_1_**) Inactive timers at the time of the CF spike lose resource; it is then taken up by the most active timer. (**G_2_**) When the resource is limited and binding kinetics are zero-order, inactive timers lose a fixed amount of resource regardless of the amount they have bound (***left***, *gray lines*, and ***right***, *gray bars*), which are then taken up by the most active timer (*red lines and bar*). (**G_3_**) When resource is plentiful and kinetics are first order, inactive timers lose resource in proportion to their current level of bound resource. (**H-K**) Different biochemical implementations lead to the eligibility window matching different features of the PF-CF interval distribution. (**H**) The single timer with broad TeMP window aligns the peak timing of the plasticity rule (i.e., the eligibility window) to the median of the distribution. (**I**) The single timer with narrow TeMP window aligns the peak timing to the nearest local maxima of the distribution (***bottom***). Arrows indicate mean direction of movement of eligibility window peak from a given initial starting point. Grey band and dark curve show 1 s.d. around the mean value of the peak eligibility as it evolves, for 30 simulations. (**J**) The multiple timer selection mechanism with resource limited, zero-order kinetics exhibits strong competition between different timers, leading to a winner-take-all operation that selects the mode of the distribution. (**K**) The proportional loss of resources in the timer selection rule with plentiful resources and first order kinetics leads to the eligibility trace approximating the shape of the distribution (shown for 22 timers).

The time at which the eligibility window opens can be adjusted by changing either the rate at which the accumulated species B* is produced or the threshold for producing the eligibility molecule C*. Temporal metaplasticity is controlled by the timing of a postsynaptic spike relative to the peak of the eligibility window (see **Fig. 4C**). In the molecular implementation, τ*_peak_* occurs close to the time at which the timer on/off switch (A_1_*/A_1_) is turned off by the eligibility molecule C* (**Fig. 5B**, *dashed vertical lines*). If the timer on/off switch is in its on state A_1_* at the time of the postsynaptic spike, then the postsynaptic spike has occurred before τ*_peak_*, so the threshold should be decreased, reducing τ*_peak_* (**Fig. 5C_1_**, ***left***). If instead the switch has returned to its off state A_1_, then the threshold should be increased, increasing τ*_peak_* (**Fig. 5C**_1_, ***right***). This scenario creates a “broad” temporal metaplasticity window covering all spike timing intervals up to some upper cutoff, because the shift in τ*_peak_* depends only upon whether the postsynaptic spike is earlier or later than τ*_peak_* (**Fig. 5C**_2_). To implement the narrow temporal metaplasticity window used in the cerebellar circuit simulation (**Fig. 4C**_3_), changes in τ*_peak_* can be limited to only occur when the concentration of the accumulated species B* is above threshold (**Fig. 5D_1_,D_2_**).

#### Biochemical implementation of multiple timer selection/combination mechanism

The multiple timer selection mechanism can be implemented with a bank of accumulation-to-threshold timers, each of whose activations is determined by the concentration of its C* molecule (**Fig. 5A,E**). Alternatively, a timer bank could be constructed from a cascade of sequentially active molecular species (**Fig. S8A**). Specifically, we assume that each timer element has a binding domain U_i_ (**Fig. 5F**, ***left***). The contribution of the i^th^ timer element to the eligibility window is then given by the fraction of its binding domains in the resource-bound state U_i_* (**Fig. 5F**, ***right***).

Changes in the eligibility window for plasticity occur due to postsynaptic (CF) spike-triggered changes in the amount of resource bound by each timer. The timer that is active at the time of the postsynaptic spike binds more resources and inactive timers lose resources (**Fig. 5G_1_**, **Fig. S8B**; cf. **Fig. 4D_3_**). When the resource is scarce and the binding kinetics are in a saturated (zero-order) regime, the change in each timer’s contribution to the eligibility window is largely independent of its current value (**Fig. 5G_2_**, *“Fixed” update rule*; **Fig. S9A,B**; **STAR Methods**). This biochemically implements the spike-based rule used in the cerebellar circuit model of **Fig. 4D_3_**. By contrast, if the resource is abundant and the binding kinetics are in an unsaturated (first-order) regime, the change in each timer’s contribution to the eligibility window is proportional to the current value of the coupling (**Fig. 5G_2_**, *“Proportional” update rule*; **Fig. S9A,C**; **STAR Methods**).

Sparse presynaptic activity, a property that has been suggested of parallel fibers^44–48^, would be particularly effective for temporal metaplasticity. This is because, for an accumulation-to-threshold molecular timer, if a presynaptic input arrives when the timer has been started, it must either reset the timer, so that the earlier presynaptic spike is effectively ignored, or be ignored itself. In fact, the single adjustable timer implementation does ignore spikes occurring immediately following a previous spike, such as would occur in bursting activity, since the timer on/off switch is already turned on. Alternatively, not every parallel fiber spike may reliably activate the timer (e.g., due to a refractory period or inhibitory gating), reducing the effective PF input rate. Or, sparse parallel fiber bursts^23,49–56^ could be the mechanism that drives metaplasticity.

These different biochemical implementations of temporal metaplasticity have strong computational implications: each implementation described above matches the eligibility window for plasticity to a distinct statistical feature of the pre-post spike-timing distribution. We illustrate this by considering a hypothetical case of a bimodal distribution (**Fig. H-K**, *gray distributions*). The broad window single-timer-adjustment rule (**Fig. 5C**) matched the median of the spike-timing distribution (**Fig. 5H**). This is because, when τ*_peak_* is at the 50^th^ percentile of the distribution, the probabilities of shifting left or shifting right are equal. The narrow window rule (**Fig. 5D**) discovers local maxima in the shape of the distribution (**Fig. 5I**). This is because the negative-then-positive shape of this window represents a temporal derivative filter that equals zero (i.e. causes no shifts) at the local maximum of the distribution. The fixed update multiple-timer-selection rule (**Fig. 5G_2_**) exhibits strong competition between timers, leading to winner-take-all behavior that selects the mode of the spike-timing distribution (**Fig. 5J**), while the proportional update rule approximates the full shape of the distribution (**Fig. 5K**; see **Supplemental information** for mathematical derivation). Functionally, the mode or local maximum rule might be optimal for a task like OKR learning with a characteristic delay. By contrast, a rule matching the full distribution might be optimal for a task in which a motor action leads to multiple feedback events arriving at a distribution of different delay times.

Overall, these models demonstrate biologically plausible mechanisms for temporal metaplasticity that are built upon simple biochemical network motifs. Each mechanism uses the same spike-timing statistics thought to drive plasticity of synaptic weights to simultaneously drive metaplasticity of the timing rules for plasticity. Different molecular implementations of these rules can tune the timing requirements for plasticity to match different features of the spike-timing distributions, suggesting there may be a whole suite of temporal metaplasticity rules in the brain that are adapted to the specific functional needs of different circuits and learning tasks.

## DISCUSSION

The timing requirements governing neural plasticity are a key component of the algorithm that a neural circuit uses to learn. The central finding of the current study is that the timing requirements for neural plasticity are not fixed, but can be dramatically altered by experience. We refer to this capacity of the brain as ***temporal metaplasticity*.**

Our results demonstrate temporal metaplasticity of short-term as well as long-term plasticity in the cerebellum. Temporal metaplasticity was induced in the physiological conditions of awake behaving animals undergoing different visual experiences in their home cages, and it was expressed in the timing rules for plasticity measured *in vitro.* Assessments of plasticity using synaptic currents (**Fig. 1C**), synaptic potentials (**Fig. 1D-E**), and synaptically-driven spiking (**Fig. 1A-B**) in Purkinje cells provided convergent evidence for temporal metaplasticity. For each measure of PF-Purkinje cell transmission, the effects of experience on the timing rules for plasticity were not subtle; experience completely reversed which pairing intervals were effective and ineffective at inducing associative plasticity.. Moreover, temporal metaplasticity was present in adulthood as well as during development.

The discovery of this new neural phenomenon opens up many exciting questions about its mechanisms and functions, which could take decades to fully elucidate, as has been the case for neural phenomena such as LTP. Our observations in the oculomotor cerebellum and our modeling results provide initial insights and evidence-based hypotheses about the mechanisms and function of temporal metaplasticity, which can guide future research.

### Candidate mechanisms of temporal metaplasticity

Our modeling work provides foundational computational principles, instantiated in biologically plausible biochemical network motifs, about how temporal metaplasticity could be induced and expressed. Notably, the modeling demonstrates that temporal metaplasticity need not require anything exotic mechanistically—it could, in principle, be instantiated with as few as four ordinary molecular elements, and controlled by the same spiking statistics that control synaptic plasticity.

#### Which spiking statistics drive temporal metaplasticity?

Our experimental results demonstrate that experience can alter the timing rules for plasticity. Although there is increasing recognition that experience can influence the brain through multiple body-to-brain signaling pathways^57^, the primary mechanism for experience to influence the brain is through the spiking activity of neurons encoding the experience. Thus, a key mechanistic question is which spiking statistics might drive temporal metaplasticity. The most straightforward hypothesis is that the same spiking statistics that control experience-dependent changes in synaptic weights, i.e., plasticity, in a given circuit, also control experience-dependent changes in the timing rules for altering weights, i.e., temporal metaplasticity. The modeling results demonstrate the plausibility of this possibility in the cerebellum, where the distribution of time intervals between parallel fiber (PF) and climbing fiber (CF) spikes could, in principle, control the induction of both LTD and the temporal metaplasticity of LTD at the PF-Purkinje cell synapses.

However, it is possible that other signals related to visual experience could influence the induction of temporal metaplasticity. Dark rearing likely induces widespread changes in the visual system and brain.^58^ This does not alter the central conclusion of the study, that the timing rules for cerebellar plasticity are not fixed, but can be altered by the inputs the cerebellum receives from other brain areas. However, it does mean that visual experience may alter not only the error signals carried by the climbing fibers, but also the other signals impinging on Purkinje cells, including parallel fiber inputs, inhibitory inputs, neuromodulatory inputs and other factors such as hormonal signaling, which could potentially contribute to the induction of temporal metaplasticity.

#### Molecular mechanism?

At the subcellular level, our models predict specific biochemical network motifs. Most notably, we predict the presence of one or more adjustable molecular timers implemented through mechanisms such as a biochemical accumulator with an adjustable accumulation rate or threshold, or a set of fixed timers activated independently or in a sequential cascade. Biologically, there are innumerable ways that such timers could be implemented and adjusted. In the cerebellum, known elements of the biochemical cascade for PF-Purkinje cell LTD are an appealing set of candidate molecular effectors of temporal metaplasticity; however, unknown molecules that modulate the dynamics of the main LTD cascade also could be elements of the molecular mechanism for temporal metaplasticity. More generally, changes in the kinetics of a molecular timer controlling plasticity might be implemented through changes in transcription, translation, post-translational modification, morphological changes, or molecular trafficking (e.g., altered localization of elements of the LTD cascade relative to each other), to name just a few. The slow time course of several weeks for temporal metaplasticity could reflect an intrinsically slow cellular process, such as protein turnover, but also could result from noise in the spiking statistics increasing the time for the temporal metaplasticity mechanism to “find” (via a random walk) a small peak in the distribution of PF-CF spike times. Given the very large space of potential molecular mechanisms, an unbiased approach may be the best initial strategy for investigating the molecular pathways for temporal metaplasticity. Importantly, there need not be a single mechanism for temporal metaplasticity, even at a single synapse, just as there is no single mechanism for LTP. Although a major challenge, success in identifying molecular mechanisms would provide the opportunity to manipulate temporal metaplasticity to test its role in circuit function and behavior, guide the search for temporal metaplasticity in other brain areas, and facilitate the evaluation of its potential role in neurological and psychiatric disorders.

### Candidate function of temporal metaplasticity

Previous studies have provided evidence for experience-dependent changes in the threshold for plasticity at a number of sites in the brain, including the PF-Purkinje cell synapses,^59–64^ and the concept of a sliding threshold for plasticity^65^ has been influential in computational studies of the function of neural networks. The *temporal metaplasticity* identified in the current study is qualitatively different from previously described *threshold metaplasticity*, not only in the way it alters plasticity (i.e., altering the preferred pairing interval rather than the amount of plasticity), but also in its potential function. Whereas threshold metaplasticity has been hypothesized to support synaptic homeostasis, synaptic tagging, memory allocation,^66^ and the general enhancement or impairment of learning by factors such as stress, arousal, an enriched environment, or exercise,^67–70^ temporal metaplasticity provides a candidate mechanism for meta-learning, or learning to learn.^71^

Our modeling explored the hypothesis that the function of temporal metaplasticity is to adaptively tune the plasticity windows to meet the functional requirements imposed by the properties of a particular circuit and behavioral learning task. More specifically, we demonstrate the biological plausibility of the idea that temporal metaplasticity in the cerebellum adaptively tunes the timing requirements for synaptic plasticity to compensate for the feedback delay for behavioral errors to be reported to the circuit, thereby improving the temporal accuracy of learning.

This hypothesis was inspired by several experimental observations, modeled in the context of the well-established relationship between signaling in the oculomotor cerebellum and behavior, which allows unusually direct comparison between cellular and behavioral observations with a resolution of a few tens of ms. First, the observed, experience-dependent timing rule for plasticity precisely matches the feedback delay for the visual error signals carried by climbing fibers. That feedback delay has been measured *in vivo* during oculomotor learning to be ∼120 ms^3,17^, and, after visual experience, associative LTD *in vitro* is induced very specifically by activating CFs at that delay relative to PFs, and not delays of 100 ms or 150 ms^3^ (**Fig 1 A-C)**.

Second, there is a striking parallel between the timing rules for LTD in the oculomotor cerebellum and the temporal accuracy of oculomotor learning in dark- and normally reared mice. The role of PF-Purkinje cell LTD in OKR learning is well established^34,72^, and changes in the firing rate of Purkinje cells in the flocculus have an approximately linear effect on the eye movement behavior.^32,39,73^ Therefore, although one cannot rule out additional effects of home cage visual experience on the OKR learning behavior, the known connections from LTD to Purkinje cell firing to eye movement make a compelling case that experience-driven changes in the timing rules for LTD could alone account for the observed changes in the temporal accuracy of learning (**Figs. 2&3**). A third experimental observation guiding the modeling was the previous finding of different rules for associative plasticity at PF-Purkinje cell synapses in different regions of cerebellum,^3,16,17^ which support different behavioral tasks. In the cerebellar flocculus, the visual feedback delay for climbing fibers to signal oculomotor errors is predictable and even conserved across species,^3,17^ hence a plasticity rule matched to this delay could, in principle, be genetically pre-specified. This pre-specification might correspond to a special case of the multiple timers model of temporal metaplasticity, with just two fixed timers and the spiking statistics encoding visual experience toggling on the expression of a plasticity rule pre-specified for 120 ms. However, if pre-specified, the advantage of having expression of the functionally relevant plasticity rule contingent on experience is not clear, nor is the advantage of retaining the capacity for the coincidence-based plasticity rule. Further, it seems less likely that plasticity rules matching the circuit requirements could be pre-specified for each of the myriad learning tasks supported by the cerebellum. For different tasks, climbing fibers carry error signals of different modalities, which could have different and almost certainly less stereotyped feedback delays than those in the oculomotor cerebellum, especially for more complex learning tasks. Thus, for the broader set of cerebellum-dependent behavioral tasks, a more general solution to the temporal credit assignment problem would be an adaptive temporal metaplasticity process that tunes the timing rules for plasticity based on the statistics of the feedback delays experienced.

Based on these considerations, the modeling demonstrates how the timing requirements for plasticity may not only be *altered* by experience, as revealed by the experimental observations, but adaptively tuned by it. Further, the modeling makes the surprising prediction that different classes of molecular mechanisms generate functionally distinct temporal metaplasticity rules that adjust the timing requirements for plasticity to different statistics of the distribution of spike-timing relationships. These different implementations could provide distinct functional advantages for different learning tasks. Thus, temporal metaplasticity could provide a general solution to the temporal credit assignment problem common to all learning circuits and thereby support a type of metalearning—an experience-dependent improvement of the temporal accuracy of learning.

Notably, appropriately tuned timing rules for associative plasticity are necessary but not sufficient for temporally precise learning. The latter also requires a suitable representation of temporal information (temporal basis set) to serve as a substrate for the plasticity rule (**Fig. S6)**. Previous studies have proposed several mechanisms for generating such neural representations of temporal information.^44,74–80^ Moreover, computational modeling has suggested that metaplasticity of nonassociative short-term plasticity (changes in the amount of synaptic facilitation or depression during a presynaptic spike train) could improve the discriminability of the neural representations of temporal information.^81^ Such a process could work in conjunction with the temporal metaplasticity of associative plasticity, described in the current study, to improve the temporal accuracy of associative learning.

### Is temporal metaplasticity a cerebellar specialization, or a brainwide capacity of synapses?

Temporal metaplasticity provides a candidate synaptic solution to the temporal credit assignment problem. This would be a powerful mechanism for optimizing almost any type of associative learning in which temporal credit assignment is necessary, including learning driven by positive or negative feedback, such as oculomotor learning, or learning the relationships between sequential events in the external world. Even for events that are simultaneous in the world, such as visual and auditory cues arising from a single object, learning the correct associations requires the alignment of neural signals with different characteristic processing times to reach a given site in the brain. Conversely, failure to adaptively define the temporal relationship between neural events whose associations should be learned could contribute to clinical conditions such as schizophrenia and autism, which are associated with an enlarged temporal window for binding events.^82^

The computational potential of temporal metaplasticity raises the question of whether this capacity is confined to cerebellar synapses. If so, it would represent a remarkable and perhaps defining specialization of the cerebellum for its hypothesized role in temporally precise learning, as exemplified by the skills of elite athletes or musicians.^83^ On the other hand, the mechanistic models of temporal metaplasticity could readily be applied to almost any heterosynaptic plasticity mechanism, including reward-driven learning, and also might be extended to homosynaptic spike-timing dependent plasticity (STDP) mechanisms controlled by the statistics of spiking in the pre- and post-synaptic neurons^84^ Likewise, challenges related to unknown propagation delays in engineered systems, such as wireless communication networks and AI systems based on recurrent neural networks^85–87^,might benefit from temporal metaplasticity-inspired solutions to the temporal credit assignment problem, based solely on local information. Temporal metaplasticity provides a potentially general and flexible mechanism for adaptively tuning the timing contingencies controlling plasticity and learning, and hence is a candidate synaptic mechanism for experience-dependent improvement in learning, i.e., meta-learning.

## METHODS

### Animals

All experimental procedures were approved by the Administrative Panel on Laboratory Animal Care at Stanford University. C57Bl/6J mice were purchased from The Jackson Laboratory (stock no. 000664). Mice used in experiments were all bred at Stanford, and housed from birth either on a reversed 12 hr light/12 hr dark cycle (normally reared mice) or on a 24 hr dark cycle (dark-reared mice) in a custom-made light-proof chamber or in a light-proof circadian chamber bought from Actimetrics (part number: PT2-CCM3). Slice physiology experiments were performed on these mice at 24-42 days of age, or behavioral experiments were performed at 8-24 weeks. Other mice were reared normally until 60 days of age, then moved to dark chambers for 60-120 days, and, in some cases, moved back to normal rearing conditions after 60 days in the dark, with slice physiology and behavioral experiments conducted at the time points indicated in the main text, **Fig. 3**, and **Fig. S5**. For mice housed in darkness, husbandry was performed using night vision goggles (Wanney Group, cat. no. WNNVE-B201A) with an additional filter (Thorlabs, cat. no. FELH0800) on the infrared emitter to minimize emission in the visible range. All animals had *ad libitum* access to food and water. Experiments on normally reared mice were conducted during the animals’ dark cycle.

### Acute slice electrophysiology

Experimenters were blind to the rearing condition during experimentation and data analysis. All reagents were obtained from Sigma-Aldrich unless otherwise indicated.

For experiments on young mice (24-42 days old), mice were anesthetized using isoflurane (1-3%) and decapitated; their brains were extracted immediately and 250 μm-thick coronal floccular slices were cut using a VT 1200 Vibratome (Leica). Brains were sliced in ice-cold artificial cerebrospinal fluid (ACSF (concentration in mM): NaCl(119), KCl(2.5), NaH_2_PO_4_(1), NaHCO_3_(26.2), MgCl_2_(1.3), CaCl_2_(2.5), D-Glucose(10)), equilibrated with Carbogen (95% O_2_, 5% CO_2_, Praxair) to maintain pH at 7.3 (osmolality ∼323 mOsm). Slices were then incubated at ∼34-36 °C for 30 mins, and then at room temperature for at least an hour.

For experiments on adult mice (60-160 days old), mice were anesthetized with avertin (2,2,2-tribromoethanol; 0.2 mL/10 g body weight; intraperitoneal injection), and then transcardially perfused with ice-cold N-methyl-D-glucamine solution (NMDG ACSF; (concentration in mM): NMDG (93), KCl(2.5), NaH_2_PO_4_(1.25), NaHCO_3_(30), HEPES(20), D-glucose(25), sodium ascorbate(5), thiourea(2), sodium pyruvate(3), N-acetyl-L-cysteine(12), MgSO_4_(10), CaCl_2_(0.5), R-CPP(0.0025) and NBQX(0.005) equilibrated with Carbogen (95% O_2_, 5% CO_2_, Praxair), with pH adjusted to 7.3 using HCl (osmolality ∼323 mOsm)) using a peristaltic pump until the liver turned pale (∼1.5 minutes). Brains were extracted immediately and 250 μm-thick coronal slices of the cerebellar flocculus were cut in ice-cold NMDG ACSF and then incubated in NMDG ACSF without R-CPP and NBQX at 32-34 °C for 10-12 min. Then the slices were transferred to room temperature ACSF and were incubated for at least an hour at room temperature.

During the electrophysiology experiments, slices were bathed in ACSF containing 100 µM picrotoxin to block GABA_A_ channels, and held at 32-35 °C. Recordings were obtained from Purkinje cells identified visually using an Olympus BX51WI upright microscope. Signals were digitized at 20 kHz using a Multiclamp 700B amplifier and low-pass filtered at 10 kHz. Data were acquired using PClamp10 and analyzed using IgorPro with NeuroMatic.

Plasticity of parallel fiber-to-Purkinje cell transmission was induced by pairing parallel fiber and climbing fiber stimulation. The climbing fiber input to a Purkinje cell was stimulated in the granule cell layer just beneath the Purkinje cell soma (∼20-50 µm from Purkinje cell layer), and its recruitment was confirmed by the presence of complex spike responses in the Purkinje cell. Parallel fibers were stimulated by placing a stimulating electrode in the upper third of the molecular layer. All stimulation was done using bipolar electrodes pulled from theta glass (Sutter Instruments).

Long-term plasticity was induced by 300 pairings of parallel fiber and climbing fiber activation at 1 Hz. Each pairing consisted of a single stimulus delivered to the parallel fibers and a single stimulus to the climbing fiber, each 150 µs in duration. The interval between parallel fiber and climbing fiber stimulation (PF-CF pairing interval) was either 0 ms (coincident stimulation) or 120 ms (parallel fibers then climbing fiber).

#### Assessment of long-term plasticity of parallel fiber-elicited spiking in Purkinje cells

Recording electrodes (3-5 MΩ) were pulled from borosilicate glass tubing (Harvard Apparatus) and filled with ACSF for extracellular recordings from Purkinje cells. Parallel fiber-elicited spiking in the Purkinje cell was tested with parallel fiber stimulation (1 pulse, 150 µs) delivered at 0.05 Hz for 5 min before, and for 30 min after the induction of plasticity. Spikes were counted in the first 50 ms (**Fig. 1**), or the first 30 or 100 ms (**Fig. S1**) after the parallel fiber stimulus and normalized to the spontaneous spike count measured in a window of 50 ms (or 30 or 100 ms for **Fig. S1**), starting 100 ms before each stimulus. Plasticity of the spike count was measured as the ratio of the normalized spike count measured at each time post-pairing divided by the normalized pre-induction spike count.

#### Assessment of long-term plasticity of parallel fiber-elicited synaptic currents in Purkinje cells

Visually identified Purkinje cells were patched and voltage clamped at −70 mV in the whole-cell configuration during testing. Series resistance was compensated. Series and input resistance changed by less than 20% throughout the recording. Cells with more than 20% fluctuations in input resistance were excluded. The recording pipette was filled with an internal solution containing, KMeSO4(133), KCl(7.4), MgCl2(0.3), HEPES(10), EGTA(0.1), MgATP(3), Na2GTP(0.3), pH adjusted to 7.3 with KOH, osmolarity 260-290 mOsm. Parallel fiber-elicited synaptic currents in the Purkinje cell were tested with parallel fiber stimulation (1 pulse, 150 µs) delivered at 0.05 Hz for 5 min before and for 30 min after the induction of plasticity. EPSCs were measured using clampfit software. Long-term plasticity was induced with 300 pairings of parallel fibers and climbing fiber stimulation, as described above, while the Purkinje cell was held in current clamp mode.

#### Assessment of short-term plasticity of parallel fiber-elicited synaptic potentials in Purkinje cells

Visually identified Purkinje cells were current clamped in whole-cell configuration by injecting a holding current, typically less than 1 nA, to hold the cell at ∼ −70 mV. Short-term plasticity was induced with a single pairing of parallel fiber stimulation (1 pulse, 150 μs) plus climbing fiber stimulation (two 150 μs pulses, 10 ms apart), as previously described.^3^ Test pulses (150 μs) were delivered to the parallel fiber 1s before and 1s after the climbing fiber stimulation to measure the change in the amplitude of the parallel fiber-elicited EPSP in the Purkinje cell induced by the single parallel fiber-climbing fiber pairing. The pairing was done with the climbing fiber stimulation delayed by 0, 20, 40, 60, 80, 100, 120 or 150 ms relative to the parallel fiber stimulation, or with climbing fiber stimulation omitted for the “PF only” control. The interval between trials (pairings) was 7s. 15 trials with the same pairing interval were delivered in a block, and blocks with different pairing intervals were delivered in a pseudorandomized order.

### Surgery

Mice were surgically prepared for behavioral experiments as previously described.^88^ In brief, while under anesthesia with 1.5-2.5% isoflurane, a custom-made head post was attached to the top of the skull using dental acrylic cement. To enable magnetic eye tracking, two stacked neodymium magnets with a total size of 0.75 x 2 mm (diameter x height, grade N50, axially magnetized, SuperMagnetMan.com) were implanted beneath the conjunctiva on the temporal side of the left eye. An angular magnetic field sensor (HMC1512, Honeywell Inc., size: 4.8 x 5.8 mm) was soldered to an 8-pin connector and attached to the skull above the eye using dental acrylic, in a plane parallel to horizontal (nasal-temporal) eye movements. Eye movements were measured by detecting changes in the magnetic field created by movements of the magnet implanted on the eye (for details see ref.^88^). Animals were allowed to recover after surgery for at least five days before behavioral experiments were performed.

### Behavioral experiments

Behavioral experiments were performed in a light-proof, sound-attenuated chamber. The head of the mouse was immobilized by attaching the head post to a custom-built restrainer. Visual stimuli were provided by an optokinetic drum that surrounded the animal and rotated about an earth-vertical axis. The drum was made of white translucent plastic with black and white vertical stripes, each of which subtended approximately 7.5° of the visual field, illuminated by shutter-controlled fiber-optic lights. To elicit the optokinetic response, the optokinetic drum was sinusoidally rotated at a frequency of 1.0 Hz, with a peak velocity of ±10°/s. Training consisted of 60 repeats of 50-second long blocks of optokinetic stimulation, with 10 seconds of rest in darkness between each block, as previously described.^37^ After each experiment, a calibration session was performed during which eye movements were simultaneously measured with the magnetic sensor and video-oculography, to determine the calibration factor for converting signals from the magnetic sensor (in volts) to eye position (in degrees).^88^

Data were analyzed in MATLAB. Horizontal eye position was computed from the voltage measured with the surgically implanted magnetic sensor. Eye position signals were filtered using a fourth-order low pass (15 Hz) Butterworth filter. Eye velocity was derived from the filtered eye position via a 10 ms-windowed Savitzky-Golay differentiation and smoothing filter. Data from stimulus cycles with a saccade or movement artifact (e.g., a blink or whisking) were excluded from the analysis using a threshold on the mean squared error of a sinusoidal fit to the eye velocity. The threshold was determined individually for each animal to optimize detection of artifacts with different levels of measurement noise. The experimenters were blind to the rearing condition of the mice while analyzing the data.

For each 50 s optokinetic reflex (OKR) adaptation training block, data remaining after artifact removal were aligned on the visual stimulus and averaged. Pre-and post-training values were obtained from the first minute and the average of the last three minutes of OKR training, respectively. A sinusoidal fit to the averaged eye velocity trace provided the measure of eye movement amplitude used to calculate the gain of the OKR (ratio of eye movement amplitude to visual (optokinetic) stimulus amplitude). The time of peak eye velocity relative to the optokinetic stimulus was extracted from the average eye velocity trace. Mice with a baseline OKR gain of less than 0.1 were excluded from further analysis (one normally reared and one dark-reared mouse). One additional dark-reared mouse was excluded from the analysis of OKR gain (**Fig. 2B**) because a reliable calibration of the magnet data using the video calibration method could not be obtained, but this mouse was included in the analysis of the timing of learned eye movements (**Fig. 2E**), for which a calibration of the eye movement amplitude was not required.

### Statistics

All statistics were performed using JMP statistical suite. A Shapiro-Wilk test was used to assess normality. Non-parametric statistical tests were used to analyze data that violated the normality assumption. For analysis of short-term plasticity, ARTool (Windows version) was used to align-and-rank transform data that were not normally distributed^89,90^ before conducting two-factorial ANOVA using JMP suite.

### Modeling of temporal metaplasticity

Past models have proposed ways for how the cerebellar circuit can overcome the temporal credit assignment problem associated with the delayed instructive feedback conveyed by climbing fibers.^91–93^ However, none of these past works predict experience-dependent changes in the timing rules for credit assignment observed here experimentally. Here we propose how temporal metaplasticity can adaptively tune these timing rules to offset the delay in instructive feedback. We propose two general mechanisms for implementing temporal metaplasticity and its activity-dependent induction based on the temporal statistics of neural activity experienced: adjustment of a single timer, and selection from a bank of timers (**Fig. 4C,D**).

We modeled synapses as receiving a sequence of a large number of parallel fiber-climbing fiber spike pairs. The interval between parallel fiber and climbing fiber spikes both drives changes in synaptic weight as a result of the synapse’s plasticity rule and simultaneously causes the eligibility window to change according to the synapse’s temporal metaplasticity rule.

To understand how each temporal metaplasticity mechanism shapes the associative plasticity rule, we modeled the spike intervals of parallel fiber and climbing fiber inputs as arising from a distribution *p_cf_*(τ) = **P**(**τ**_cf_ = τ), defined between τ_min_ = 0 ms and τ_max_ = 200 ms, where τ_c_*_f_* = *t_cf_* − *t_pf_*, i.e. equals the interval between the climbing fiber-driven postsynaptic complex spike time *t*_cf_ and the preceding presynaptic parallel fiber spike time. This distribution is a property of the cerebellar circuit and its inputs, but has not been measured experimentally due to the challenges of recording the spiking of cerebellar granule cells. We used either a distribution (**Fig. 4I**) derived empirically within a simulated cerebellar circuit model with simultaneous plasticity and temporal metaplasticity (**Fig. 4E**, **Supplementary Information**) or considered an abstract distribution that enabled us to highlight features of different implementations of the temporal metaplasticity rule. The abstract distribution took the form of a bimodal distribution with differing heights of peaks given by density function

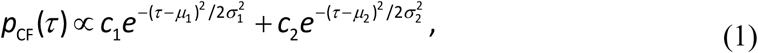

with μ_1_ = 50 ms, μ_2_ = 140 ms, σ_1_ = 50 ms and σ_2_ = 10 ms, and *c*_1_ = 1, *c*_2_ = 2 (**Fig. 5H-K)**. This distribution allows us to clearly visualize the distinct features of the distribution selected by the different temporal metaplasticity mechanisms.

Below, we describe each class of temporal metaplasticity mechanism, first at the level of a spike-timing based rule and then in terms of example biochemical network implementations.

#### Single timer adjustment mechanism

For the single timer adjustment mechanism (**Fig. 4C**), each synapse contains a single molecular timer, started by the parallel fiber, that determines the window of eligibility (**Fig. 4C_2_**, *e*(τ)) for the synapse to undergo climbing fiber-driven plasticity. The timer can be modeled as having a single variable, for example the time of the peak eligibility with respect to PF input, τ_peak_, that is adjusted by the statistics of PF-CF spike intervals. If a climbing fiber spike occurs before the eligibility has reached its peak, then the timer is too slow, and in this mechanism, the timer is adjusted so that eligibility peaks earlier. Conversely, if the climbing fiber spike occurs after the peak eligibility, then the timer is too fast and is adjusted so that the eligibility peaks later.

Mathematically, we can represent this as

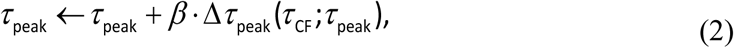

where β is a learning rate and *Δ*τ_peak_(τ_peak_; τ_CF_) is the temporal metaplasticity window, a function of the PF-CF interval and the current peak eligibility time, which determines the adjustment to the peak timing (**Fig. 4C_3_**).

*Timer adjustment with broad temporal metaplasticity window*.

The simplest version of the rule is

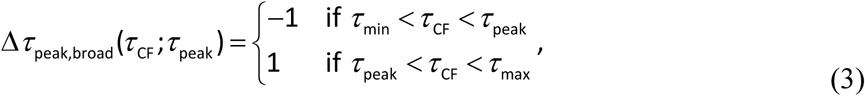

which we refer to as the “broad” temporal metaplasticity window (cf. **Fig. 5C**). The broad timer adjustment mechanism shifts the eligibility for plasticity so that its peak is at the median of the distribution of intervals between parallel fiber and climbing fiber spikes (**Fig. 5H**; see **Supplemental Information** for derivation).

We implemented this mechanism biochemically by constructing an accumulation-to-threshold timer,^94–96^ for which a reaction diagram is shown in **Fig. 5A**. In the following, we use the convention that X^∗^ refers to an activated form of a given species X, and we assume that concentrations are normalized to a fixed total for each species (i.e., X^∗^ + X = 1). Parallel fiber input causes a fast buildup of 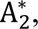 turning on a bistable molecular switch in which A_1_is quickly and maximally activated.^97^ 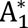 is required to produce a molecular species B^∗^that builds up in the synapse. Once species B^∗^reaches a (soft) threshold,^97^ species C^∗^ is produced, which controls the eligibility (i.e., the time-varying concentration of species C^∗^ defines the spike-timing dependent plasticity eligibility window). Species C^∗^ causes the degradation of element 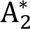, which returns 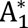 to its inactivated form. This in turn leads to the inactivation of species B^∗^and then species C^∗^, closing the eligibility window (**Fig. 5B**). Each of the four species can exist in an active or inactive form, with total concentration of each species conserved by all reactions. The dynamical evolution of the fractional concentrations (relative to the total concentration of active and inactive forms) of these species is given by

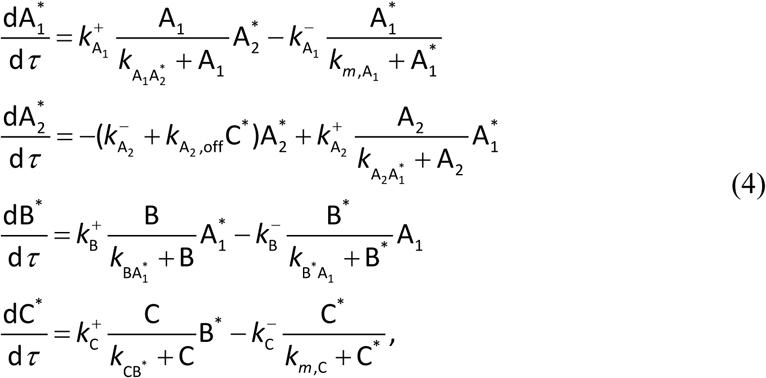

where asterisks indicate the active form of each species. Parameter values are listed in **Table S1**.

The peak timing of the eligibility can be adjusted by changing the speed at which B^∗^reaches threshold (**Fig. 5B**). Here, because the threshold is effectively controlled by the ratio of the rates of activation and inactivation of C^∗^, 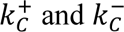, we model the rates as 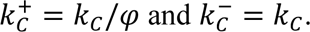. Alternatively, the peak timing of the eligibility could be adjusted by changing the rate at which B^∗^is built up. The temporal metaplasticity window is constructed as a combination of the amount of active and inactive *A*_1_present at the time of CF spike,

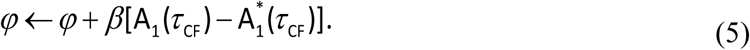

(**Fig. 5C_2_**). This could be implemented biologically by modeling the binding of B^∗^ and C as requiring another factor Y, whose concentration is changed during the time of a climbing fiber input by an amount proportional to (Y^∗^)^2^ (i.e., inactivation by dimerization).

We simulated this rule for the bimodal distribution described above. Starting from an initial value of φ = 0.01, corresponding to a steep initial rise of the activated eligibility molecule C* with a peak of around 10 ms, we calculated the change in the eligibility window due to a sequence of 200,000 PF-CF pairing intervals drawn from the distribution, for 30 independent simulations. We set the rate of temporal metaplasticity to β = 10^−3^. In **Fig. 5H**, we plotted the rule using the average value of φ over the last 10,000 intervals (sampled every 100), across all the simulations.

#### Timer adjustment with narrow metaplasticity window

We also consider a form of the timer adjustment mechanism in which changes to the eligibility window are driven mainly by PF-CF intervals that are temporally close to the current peak of eligibility,

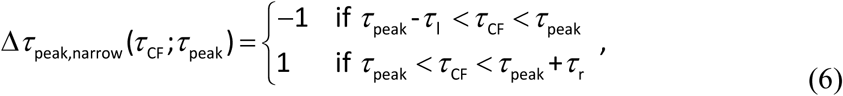

where τ_*l*_ and τ_*r*_ define the left and right sides of a narrow window around the peak. This temporal metaplasticity rule drives the eligibility window to move according to an approximate derivative of the PF-CF interval distribution, and reaches steady state near a local maximum of the distribution (**Fig. 4J**).

This temporal metaplasticity rule can be implemented using the same molecular timer described above for the broad window, Eq. (4). The temporal metaplasticity window is constructed as

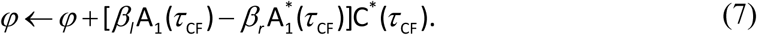

where β_*l*_ and β_*r*_ give the amplitudes of the left and right lobes of the temporal metaplasticity window. We note that, with this implementation, the left and right lobes of the temporal metaplasticity window are not symmetrical, so the resulting eligibility window may not center exactly on the peak of the PF-CF interval distribution (see ***Returning to a default timing in the dark*** below). In the simulation of **Fig. 5I** we therefore increased the amplitude of the right lobe relative to the left lobe. We simulated this rule for the bimodal distribution in the same manner and with the same parameters as for the broad window implementation. In **Fig. 5I (*top*)**, we plotted the rule using the average value of φφ over the last 10,000 intervals (sampled every 100), across all the simulations. In **Fig. 5I (*bottom*)**, we plotted a band representing one standard deviation around the mean trajectory across simulations of the peak of the eligibility window, τ_peak_.

#### Multiple timer selection mechanism

For the timer selection mechanism (**Fig. 4D**), the eligibility window for plasticity is determined by a weighted sum of activations {C^∗^ } of a bank of *M* molecular timers, with eligibility coupling strengths {*u*_*m*_},

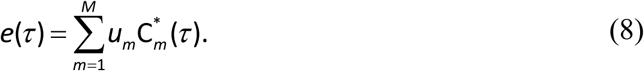

The time course of eligibility and hence the timing requirements for plasticity are altered by changing the coupling strengths {*u*_*m*_}. The selection mechanism itself is quite general, and does not depend on the particular implementation of the individual timers, so long as the bank of timers are started by arrival of the parallel fiber input and generate peaks that approximately tile time (**Fig. 4D_1_**, **D_2_**). We first discuss the rules for updating the coupling strengths, then discuss our particular implementations of a bank of accumulation-to-threshold timers and a biochemical cascade model.

#### Temporal metaplasticity rules for timer selection

We consider two methods for updating the eligibility coupling strengths {*u*_*m*_} following the arrival of each climbing fiber input, termed “additive” (winner-take-all) and “proportional,” as detailed below. For both methods, eligibility coupling strengths are increased or decreased additively following the climbing fiber spike,

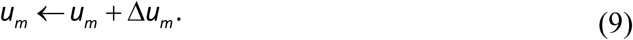

If the timer activations each have the same integrated area, both methods keep the total integrated area under the eligibility trace constant by increasing the eligibility coupling strength *u*_*m*_ of the timer that is most active at the time of the climbing fiber input and decreasing the eligibility coupling strengths of the other timers.

For the fixed update method, a fixed amount δ > 0 (up to thresholding to keep the coupling strengths non-negative) is subtracted from the eligibility coupling strengths of all the timers except the one (timer *n*) that is most activated at the time of the climbing fiber input,

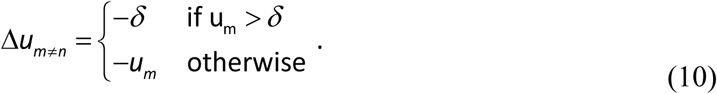

The coupling of the most activated timer is then increased by an amount

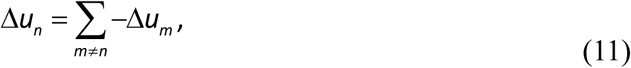

so that the sum total coupling change is 0. As discussed in the main text, this could arise biochemically as a consequence of the binding kinetics of a limited resource that controls the coupling of the timers to the eligibility for plasticity: these limited resources unbind at the time of the climbing fiber spike from out-of-sync timers and are transferred to the most in-sync timer (**Fig. 5F,G_1_,G_2_**; **Fig. S8B**, **S9**; see **Supplemental Information** for details). We note that alternative forms of competition like winning timers actively suppressing losing timers could also be implemented. These additive updates result in winner-take-all dynamics, as out-of-sync timers have their coupling to eligibility reduced to zero (**Fig. 4K**, **5J**). For an idealized set of timer activations, and assuming that the PF-CF interval distribution has a unique mode, we show that this results in an eligibility window that consists only of the timer that is active when the mode interval occurs (**Supplemental Information**).

For the proportional update method, the eligibility coupling strength *uu*_*mm*_ of the timer that is most active at the time of the climbing fiber spike is increased by an amount proportional to its difference from the maximal coupling strength (here, defined to be 1), while decreasing the coupling of the other timers by an amount proportional to their current coupling strength,

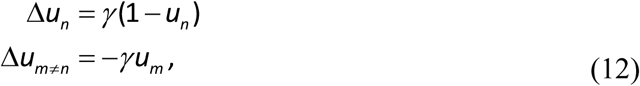

where γ > 0 is a learning rate. This update automatically preserves the sum of the eligibility coupling strengths and assures that all coupling strengths remain within the range [0, 1]. The proportional coupling update method can be biochemically implemented in a manner similar to the winner-take-all method (**Fig. 5G_3_**, **Fig. S9**; see **Supplemental Information** for details). For ease of simulation, for all simulation results shown in **Figs. 4** and **5** we used the instantaneous updates described in Eqs. (10)–(12). For an idealized set of molecular timers with identical, narrowly peaked, symmetrical activations that tile time and a stationary distribution of intervals between parallel fiber and climbing fiber spikes, the proportional update rule results in an eligibility window that can be shown (see **Supplemental Information**) to approximate the shape of the distribution (cf. **Fig. 5K**).

#### Biochemical implementation of molecular timer bank

The bank of molecular timers can be constructed in at least two ways. First, the timer bank could be a set of accumulation-to-threshold timers with distinct characteristic values of τ_peak_ (**Fig. 5E**). These might exist within a single synapse, or each of the ∼100,000 parallel fiber-Purkinje cell synapses could contain a single timer with a different timing. Second, the timers in the bank could arise from a cascade of sequentially activated molecules, each of which is activated by the previous element in the chain, with the first element corresponding to the parallel fiber input (**Fig. S8A**). Each of these particular implementations fall into the same general class of mechanism in which the eligibility trace is built up from a temporal basis set defined by a bank of timers and their eligibility coupling weights.

We simulated temporal metaplasticity using a bank of accumulation-to-threshold timers with either the fixed or proportional update rule, and which experienced PF-CF intervals chosen from the bimodal distribution. For the fixed update rule, we used a bank of *M* = 11 timers, with values of φφ chosen so that the peaks of the timers tiled 200 ms. At the start of the simulation, the coupling strength (which can be mapped to concentrations of bound resource molecules; see **Supplemental information** Equation S.8) was 1 for the timer peaking closest to 0, and 0 for all the others. We calculated the change in the eligibility window due to a sequence of 100,000 PF-CF pairing intervals drawn from the distribution. The rate of temporal metaplasticity was set by δ = 10^−4^ (see Equation 12). In **Fig. 5J**, we plotted the rule using the coupling strength at the end of the simulation. For the proportional update rule, the simulation was performed in the same way, except that we used a bank of *M* = 22 timers with evenly spaced peaks to better visualize the relationship between the final eligibility window and the shape of the distribution in **Fig. 5K**.

#### Returning to a default timing in the dark

Our experimental results show that, in the dark, the timing preference of the associative plasticity rule changes to a value close to coincidence of parallel fiber and climbing fiber inputs (**Figs. 1, 3**). One possibility is that the PF-CF interval distribution has a peak at such a time, driven by nonvisual climbing fiber signals. Alternatively, it is possible that there is no peak timing (i.e., a uniform interval distribution). Here we propose modifications to the narrow window timer adjustment mechanism and the winner-take-all timer selection mechanisms such that, for a uniform interval distribution, they both tune to a characteristic delay in the light and approach a near-coincident default timing in the dark.

#### Asymmetric narrow window timer adjustment

For the narrow window timer adjustment mechanism, we note that a tendency to go to coincident timing can be achieved for a uniform distribution by increasing either the width (τ_*ll*_) or the amplitude (β_*ll*_) of the left (before the peak) lobe of the metaplasticity window, relative to the right lobe (**Fig. S7A**, ***top***). If the difference in the widths of the lobes is small, then the rule will still function approximately as a local peak detector in the light, with a small bias toward zero (**Fig. S7A**, ***middle***) and return to a close-to-coincident timing in the dark (**Fig. S7A**, ***bottom***). In addition, to enhance the speed of finding the 120 ms peak in the light, it is helpful if the rule includes a small broad window component so that the rule does not have to random walk from a near-coincident initial zero in the face of the small leftward bias inherent to the asymmetric narrow portion of the window. We note that our biochemical implementation of this rule has an asymmetry that would produce this behavior.

#### Biased winner-take-all timer selection

We modified the winner-take-all timer selection mechanism by introducing a slight bias in the coupling strength update for the timer with peak activity closest to coincidence, C_1_. For each PF-CF pair, if C_1_ is inactive, instead of decreasing its coupling strength by δ, its coupling strength is decreased by a smaller value δδ_1_ < δ (**Fig. S7B**, ***top***). In the light, as long as the distribution has a characteristic timing such that the probability of activating the mode timer is larger than the probability of activating any other timer by more than the difference between δδ_1_and δδ, the rule will still correctly align the eligibility to the peak timing (**S7B**, ***middle***). For a uniform distribution in the dark, the rule will slowly drive the eligibility window toward a coincident default (**Fig. S7B**, ***bottom***).

#### Simulation details

All simulations were run in Python 3.13, using the SciPy ecosystem (NumPy, SciPy, Matplotlib, and Jupyter/IPython). Differential equations were solved using the BDF method solve_ivp in the SciPy ODE integration package).

## Code availability

All code is available on GitHub.

## Supporting information

Supplementary Information is available for this paper.

## Acknowledgements

We thank Maxwell Gagnon and Brian R.P. Angeles for help with codes for data acquisition and analysis, and Pragna Gaddam, Macarena Martínez Rey, Danie Geimer, Paul Gorka and Jaydev Bhateja for technical assistance. We also thank James Dang and Dr. Dong Cheol Jang for help with running OKR experiments. We also thank Benjamin Lankow, Alireza Alemi, and members of the Raymond and Goldman labs for helpful discussions. **Funding:** Supported by NIH R01 DC04154 (J.L.R.), R01 NS072406 (J.L.R.), R01 NS104926 (M.S.G.), the Simons Collaboration on the Global Brain 543031 (J.L.R.) and 542989 (M.S.G.), a Stanford School of Medicine Dean’s Fellowship and a Wu Tsai Neurosciences Institute Interdisciplinary Scholar Award (S.J.), and a Stanford Center for Mind, Brain, Computation and Technology Traineeship and a Stanford Interdisciplinary Graduate Fellowship (B.J.B.).

## Author contributions

S.J., A.S., and J.L.R. designed the experiments; S.J., A.S., J.D., and M.K. performed the experiments and analyzed the data; B.J.B., M.S.G., and J.L.R. designed the modeling study with input from S.J.; B.J.B. and M.S.G. ran the simulations and performed mathematical analyses; and S.J., B.J.B., M.S.G., and J.L.R. made the figures and wrote the manuscript with input from all authors.

## Data and materials availability

All the data used to support the conclusions of the paper are presented within the paper and its supplementary materials, and are available upon request.

---

**Figure S1.**
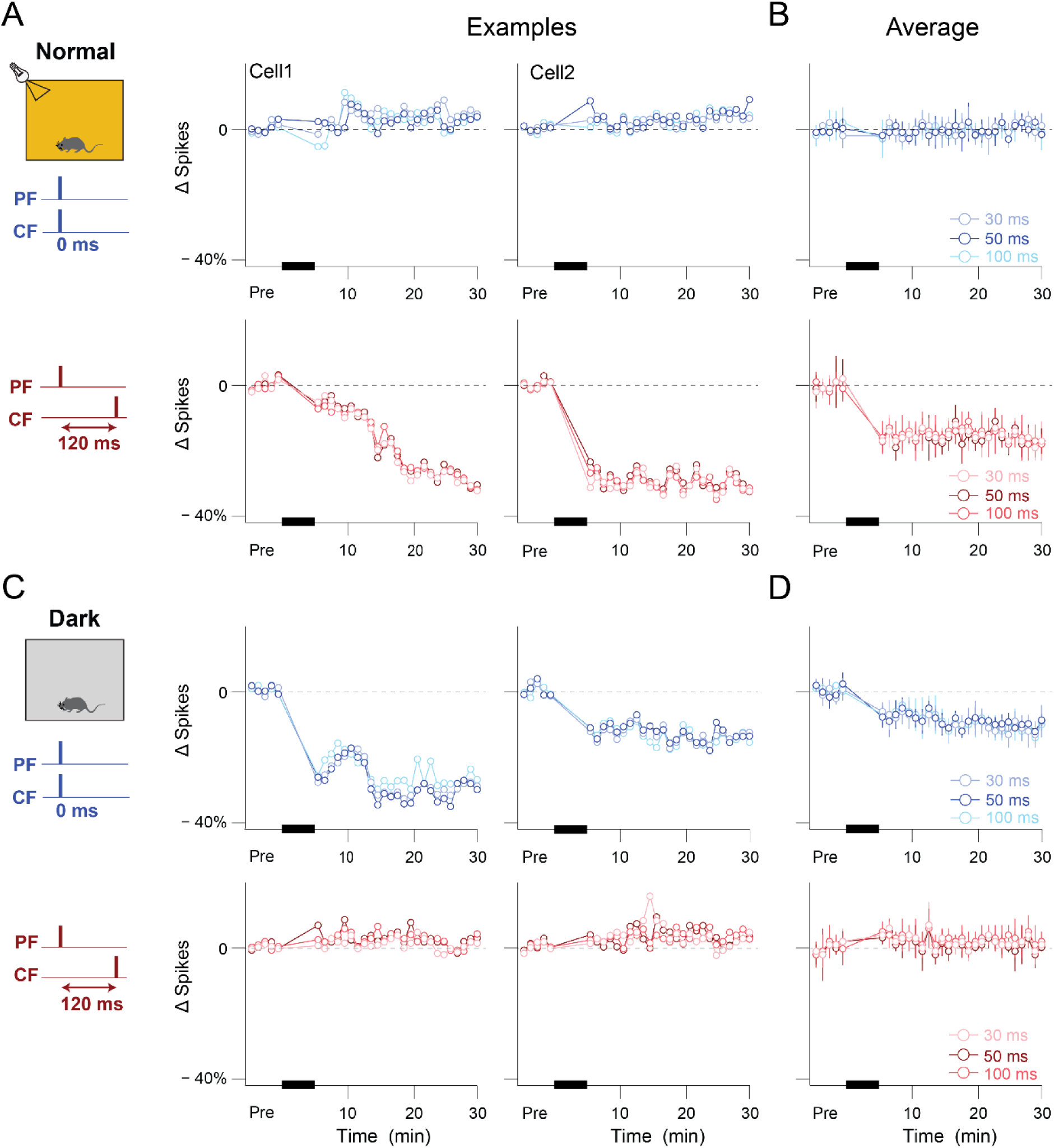
Similar plasticity of Purkinje cell spiking during the first 30, 50, and 100 ms after a parallel fiber stimulus. (**A,B**) Plasticity of parallel fiber-elicited spiking in representative Purkinje cells (**A**) and the population averages (**B**) from normally reared mice, with spikes counted in the 30, 50 or 100 ms after parallel fiber (PF) stimulation. The spike counts in these different measurement windows underwent similar changes after pairing parallel fiber (PF) and climbing fiber (CF) stimulation (300x at 1 Hz, *blue/red bars*), at pairing intervals of 0 ms (*blue; **top***) or 120 ms (*red*; ***bottom***). Results obtained using the 50 ms window for measuring spike count (*medium shade of red or blue*) are reproduced from **Fig. 1**. (**C,D**) Same as in (**A,B**), but for dark-reared mice.

**Figure S2.**
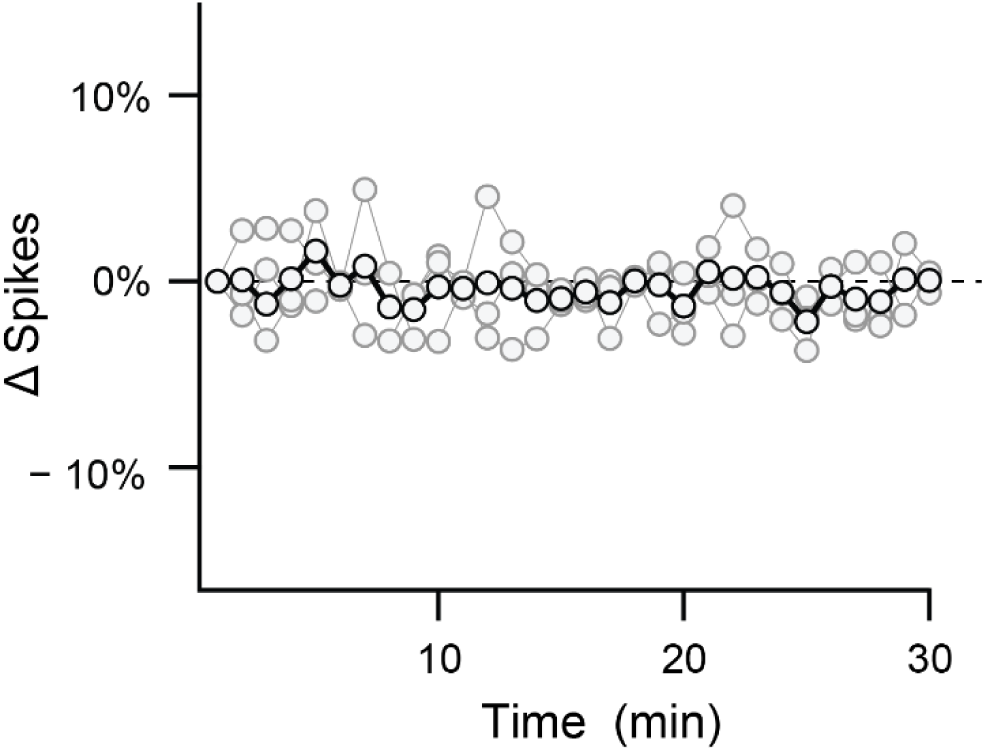
Parallel fiber-elicited spiking in Purkinje cells is stable. Parallel fiber-elicited spiking in Purkinje cells was tested at 0.05 Hz in slices of the cerebellar flocculus in the absence of any PF-CF pairing, and was found to be stable. *Grey*, individual cells, *Black*, average.

**Figure S3.**
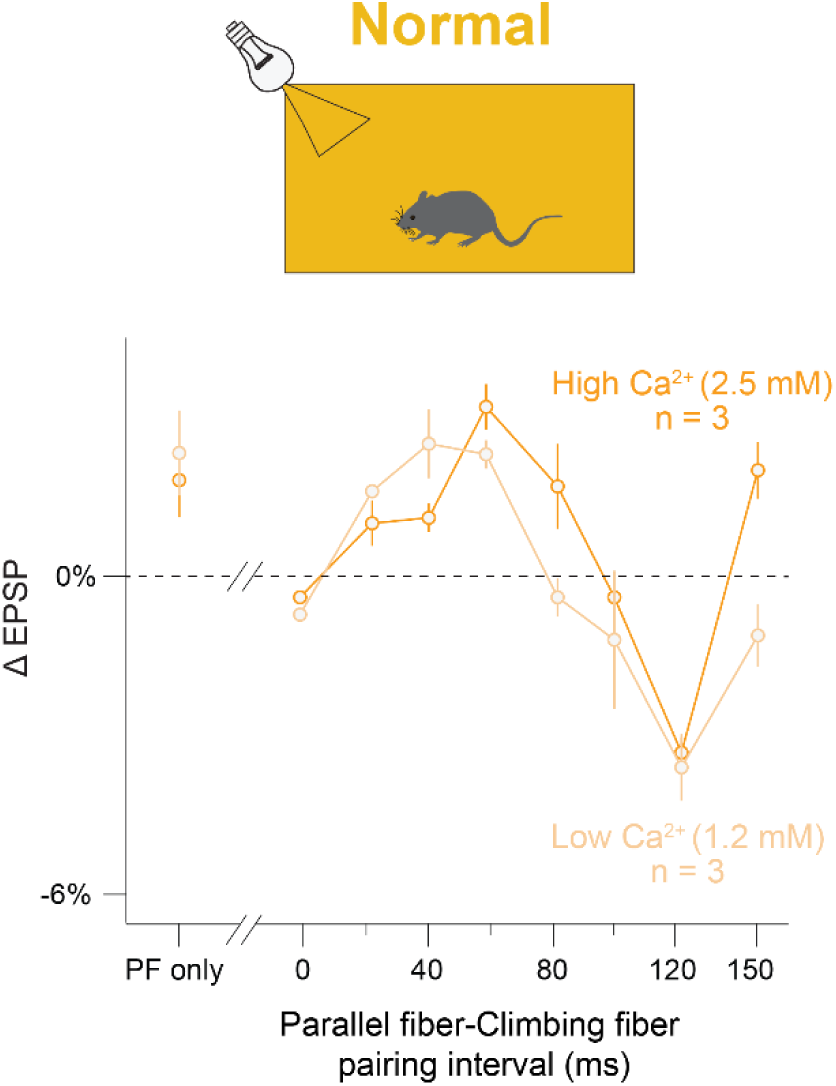
The timing contingencies for plasticity at the parallel fiber-Purkinje cells synapses were robust to extracellular calcium concentration. Short-term plasticity of PF-elicited synaptic potentials (EPSPs) in Purkinje cells, induced by a single pairing of parallel fiber (PF) and climbing fiber (CF) stimulation, was tested in slices of the flocculus of normally reared mice. The short-term plasticity induced by a range of PF-CF pairing intervals was first tested with a Ca^2+^ concentration of 1.2mM in the bathing solution (*light gold*), then the Ca^2+^ concentration in the bathing solution was increased to 2.5mM and the short-term plasticity induced at the same synapses by the same PF-CF pairing intervals was retested (*dark gold*). The calcium concentration did not alter timing contingencies for plasticity (P ≃ 1, interaction effect for [Ca^2+^] x pairing interval; P < 0.0001, main effect of pairing interval; P ≃ 1, main effect of [Ca^2+^]; 2-factor ANOVA following align-and-rank transformation of data using ARTool).

**Figure S4.**
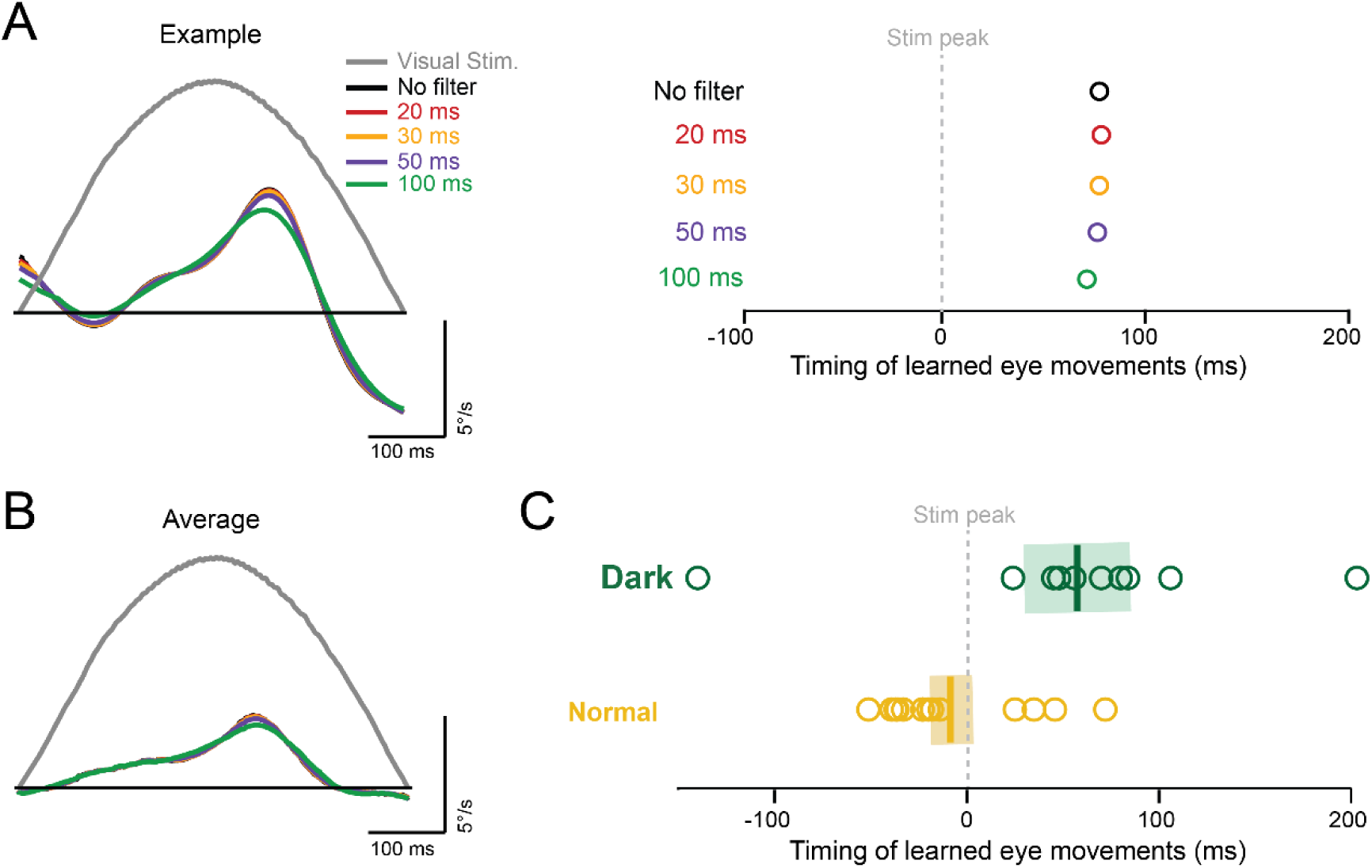
Measured differences in the timing of learned eye movements in normal and dark-reared mice are robust to temporal filtering of the eye velocity data. The values for the timing of the learned eye movements plotted in **Figs. 2E** and **3A** were measured by calculating the learned eye velocity at each time during the optokinetic stimulus, as described in the Methods and illustrated in **Fig. 2C**, and then reporting the time of the peak learned eye velocity. To assess the robustness of this measure of the timing of learned eye movements, we compared the values calculated in this manner (in which the eye movement responses were only filtered by the 10 ms filter used during differentiation of eye position to obtain eye velocity; see Methods), with the values obtained after additional temporal filtering of the traces of learned eye velocity. (A) *Left,* The learned component of the eye velocity response from one individual dark-reared mouse with no additional filtering (*black*) and after filtering with a sliding window of 20, 30, 50, or 100 ms (*colors*). *Right*, Time of the peak learned eye movement with respect to the peak optokinetic stimulus velocity for the representative mouse, calculated using different amounts of temporal filtering. **(B)** Same as in **a**, but for the learned component of the eye movements, averaged across dark-reared mice (N = 8 mice). **(C)** Timing of the peak learned eye movements (relative to the peak velocity of the optokinetic stimulus) calculated using a 100 ms sliding window average to filter the learned eye movement response in each dark-reared (*green circles)* and normally reared (*gold circles*) mouse. The learned eye movements were delayed in the dark reared mice compared to normally reared mice (*\*\*P* = 0.0065, Wilcoxon rank sum test), as also observed when the timing was calculated without the additional filter (**Fig. 2E**). *Solid vertical lines*: average peak learned timing across animals; *shaded region*: ± S.E.M.

**Figure. S5.**
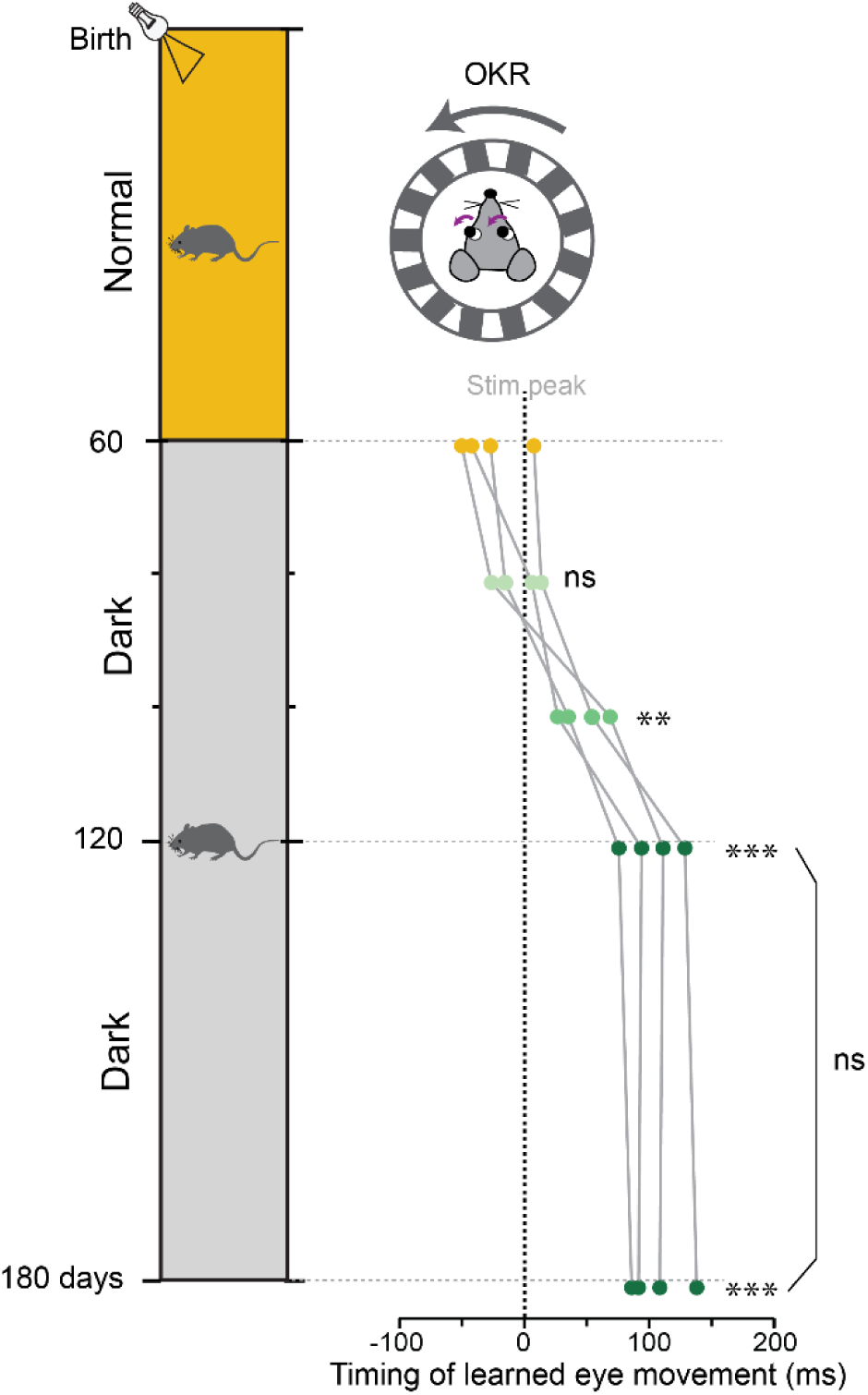
Effect of prolonged dark experience on the temporal accuracy of learning plateaus after approximately 60 days. ***Left,*** Schematic showing the timeline of manipulating the visual experience in adulthood. Mice were normally reared on a 12 hr light/12 hr dark cycle until 60 days post-birth, then maintained in prolonged darkness for the next 120 days (postnatal days P60-P180). ***Right,*** OKR learning was tested just before and 20, 40, 60, and 120 days after the mice were moved to dark housing. The timing of the learned eye movements (relative to peak velocity of the optokinetic stimulus) are plotted as in **Fig. 3A**. During the first 60 days in darkness, there was a progressive delay in the timing of the learned eye movements (*shades of green;* F_4,24_ = 24.22, *P* = <0.0001, ANOVA; P60 vs P100, \*\**P* = 0.0005; P60 vs P120, \*\*\**P* < 0.0001, Tukey). An additional 60 days in darkness induced no additional change in the timing of the learned eye movements (P120 vs P180, *P* = 0.98; P60 vs P180, \*\*\**P* < 0.0001; Tukey).

**Figure S6.**
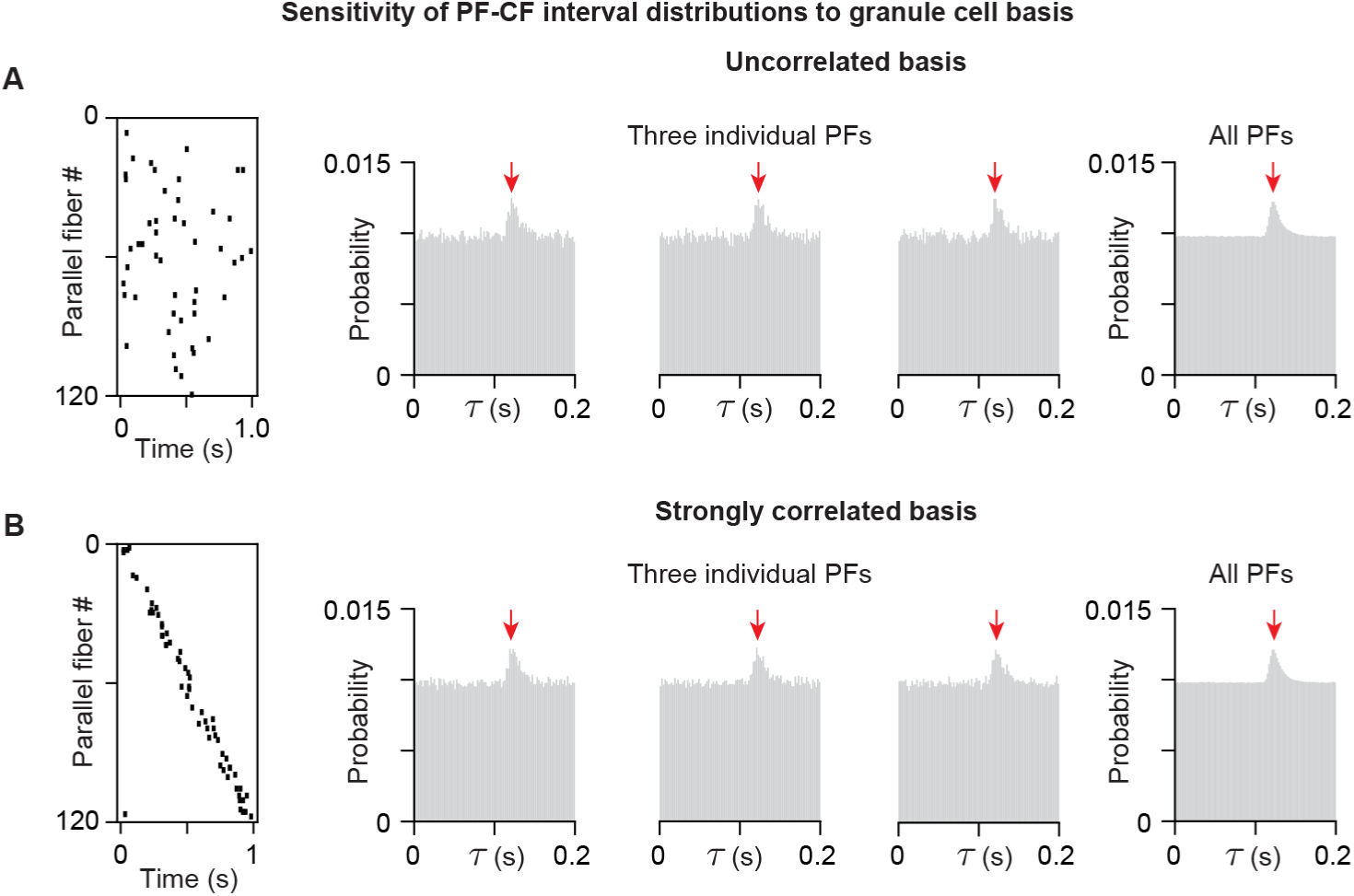
Parallel fiber (PF)-climbing fiber (CF) interval distributions, which determine the timing of the eligibility window through temporal metaplasticity, are robust to choice of PF input basis. (A) Uncorrelated parallel fiber basis. ***Left***, the uncorrelated parallel fiber spiking basis used for parallel fiber input in the home cage in the cerebellar circuit model of the main text (**Fig. 4E-K**). ***Right***, example PF-CF interval distributions for three individual PFs and for the average of all PF-CF interval distributions. All show a prominent peak close to 120 ms, characteristic of the feedback delay in the circuit. (B) Strongly correlated parallel fiber basis. ***Left***, a strongly correlated parallel fiber spiking basis, taken from the basis used during the brief period of OKR adaptation at the start and end of the simulation of the cerebellar circuit model of the main text (**Fig. 4E-K**). ***Right***, if we additionally use this strongly correlated parallel fiber spiking basis to model the parallel fiber activity during the time in the home cage, the resulting PF-CF interval distributions again show the prominent peak close to 120 ms that is characteristic of the feedback delay in the circuit. As a result, running the TeMP rules with this strongly correlated basis leads to nearly identical alignment of the timing rules for plasticity as shown in the main text (**Fig. 4J, K**).

**Figure S7.**
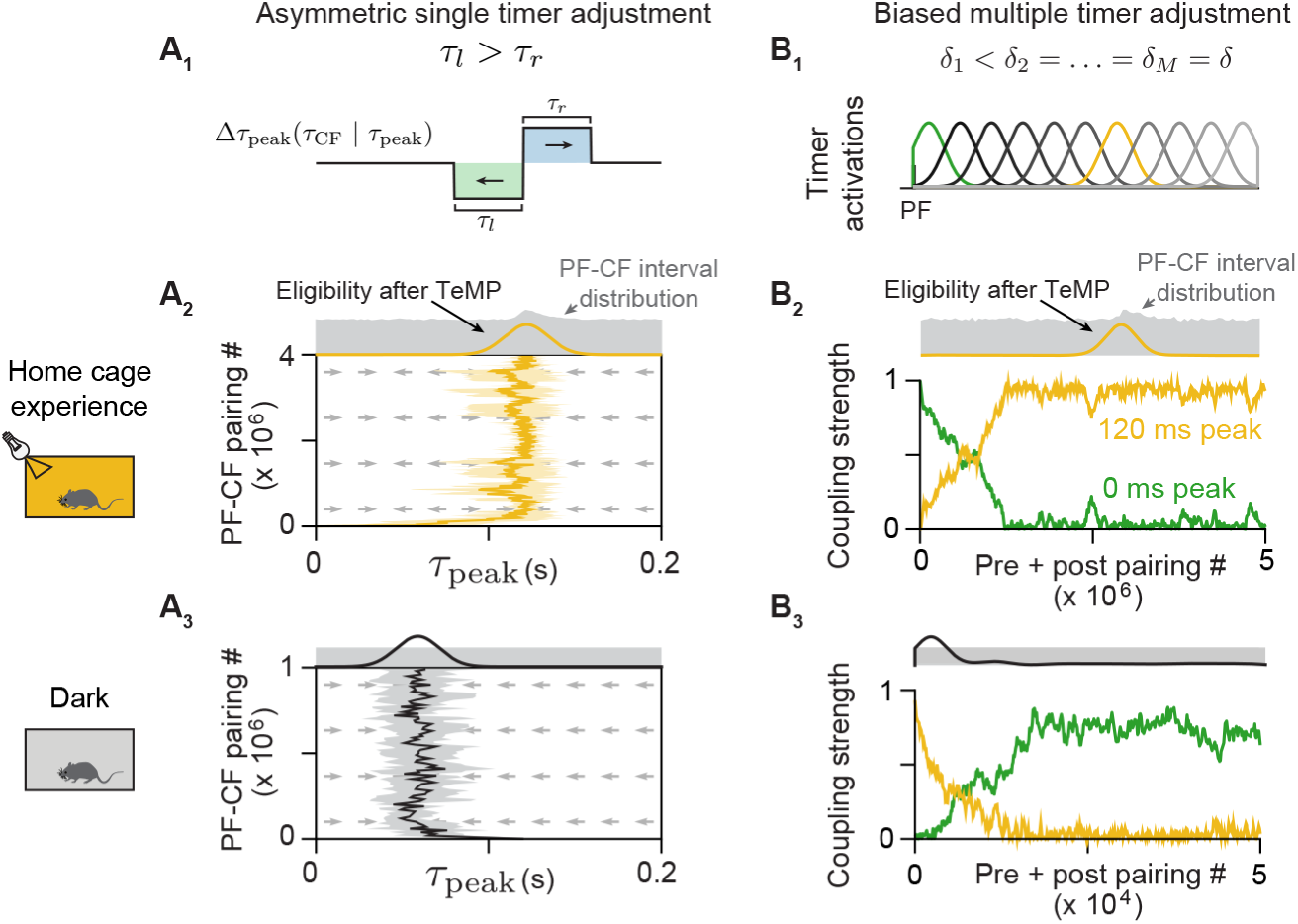
Temporal metaplasticity can reproduce the observation that the timing requirements for plasticity revert to small delays in the dark, even in the absence of structured correlations in the parallel fiber-climbing fiber spiking relationships. In the dark, we experimentally observed that the timing requirements for plasticity reverted to a preference for parallel fiber and climbing fiber spike intervals at small delays. This could be accomplished in several ways. For example, there could be a default mechanism entirely unrelated to pairing intervals that reverts the timing to small delays in the absence of visual input. For mechanisms that are related to pairing intervals, first, the parallel fiber-climbing (PF-CF) fiber spike interval distribution in the dark may have a peak at small delays, as might occur if there are fast internal feedback loops between Purkinje cells and climbing fibers. If so, the eligibility trace could be tuned to this peak in the same manner as was shown in the main text for normal visual experience (**Fig. 4J, K**). Second, as demonstrated here, even in the absence of such a default mechanism or peaks at small delays in the PF-CF pairing interval distribution, temporal metaplasticity can tune the eligibility window to small delays. This is shown for the two ‘peak-detecting’ mechanisms that were shown in the main text to be effective for oculomotor plasticity: the single timer adjustment mechanism with narrow temporal metaplasticity (TeMP) window (panels **A_1_-A_3_;** cf. **Fig. 4C, J**) and the winner-take-all version of the multiple timer selection mechanism (panels **B_1_-B_3_;** cf. **Fig. 4D, K**). (A) Single timer adjustment mechanism with slightly asymmetric, narrow TeMP window plus a very small broad component (**STAR Methods**) moves the eligibility trace to small delays when PF-CF spiking intervals are unstructured. (**A_1_)** Temporal metaplasticity window. (**A_2_) *Top***, for the PF-CF interval distribution (*grey shading*) found in the cerebellar circuit simulation (**Fig. 4E-K**) with a peak at 120 ms, the eligibility window (*black curve*) consistently finds this peak when starting from an initial value at small delays. ***Bottom***, flow fields defining the average drift from any starting τ_peak_ location (*arrows*) and 1 s.d. (*light yellow*) around the mean (*gold*) trajectory of 10 simulated trajectories corresponding to 500,000 PF-CF pairings drawn from the PF-CF interval distribution. (**A_3_**) For a PF-CF interval distribution with no peak, the asymmetry in the TeMP window drives the eligibility trace to a final peak value at small delays. ***Bottom*** panel shows 1 s.d. (*gray*) around the mean (*black*) trajectory. In this simulation, the final non-coincident timing, and corresponding rightward arrows near time zero, reflect boundary effects from the model’s simplifying assumption that temporal metaplasticity cannot be induced at negative PF-CF pairing intervals. (B) Biased winner-take-all, multiple timer selection mechanism moves the eligibility trace to small delays when the PF-CF pairing intervals are unstructured in the dark. (**B_1_**) We biased the winner-take-all competition by slightly decreasing the amount δδ_1_ by which the timer with peak activation at a PF-CF interval close to 0 ms (*green*) decreases its coupling strength u_1_ (see **Fig. 4D_3_**, “TeMP rule”) when it is out-of-sync with CF input). The timer with peak activation closest to 120 ms is indicated in *gold*. (**B_2_**) For a PF-CF interval distribution with peak at 120 ms (***top***), the coupling strength of the timer with activation closest to this peak (*gold*) grows over time while that of the 0 ms (*green*) and other timers decrease to zero, resulting in an eligibility trace peak at approximately 120 ms. (**B_3_**) Same as middle for a PF-CF interval distribution with no peak. The asymmetry in the δδ values makes the timer with peak activation close to 0 “win,” resulting in an eligibility peak close to 0 ms.

**Figure S8.**
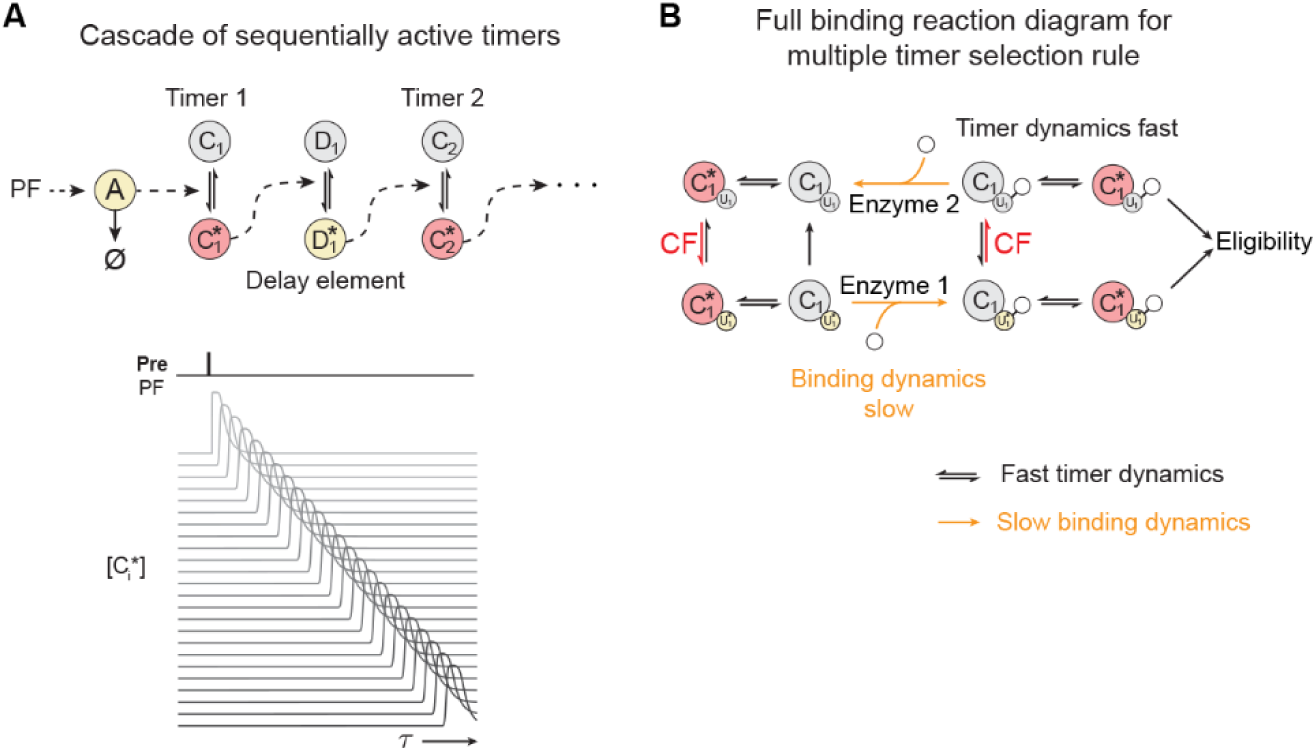
Additional reaction diagrams for the timer bank and coupling to eligibility of the multiple timer selection models. (A) Implementation of the timer bank of the multiple timer selection models through a delay line of sequentially active/activated timers. ***Top***, PF input starts the cascade through a “timer on” molecule A that rapidly degrades (indicated by “ø”) following activation. Activation of A then triggers the sequential, transient activation of the timer species C_i_ and delay elements D_i_. ***Bottom***, simulation of the sequential activation of the timer elements following the presynaptic (PF) spike. (B) Full binding reaction diagram describing how a given timer C_1_ and its binding domain u_1_ change between all of their possible inactive, active, and resource-bound forms. Unlike the simplified diagram of the main text (**Fig. 5F**), in the full reaction diagram, the binding domain can separately be active without being bound to resource and can be inactive while being bound to resource. These states occur only briefly or are irrelevant to creating the eligibility trace because they are not involved in changing coupling strengths, and therefore we collapse the four possible states of binding domain activation and binding of resource into two states of “binding domain activated and bound” and “binding domain inactive and unbound”.

**Figure S9.**
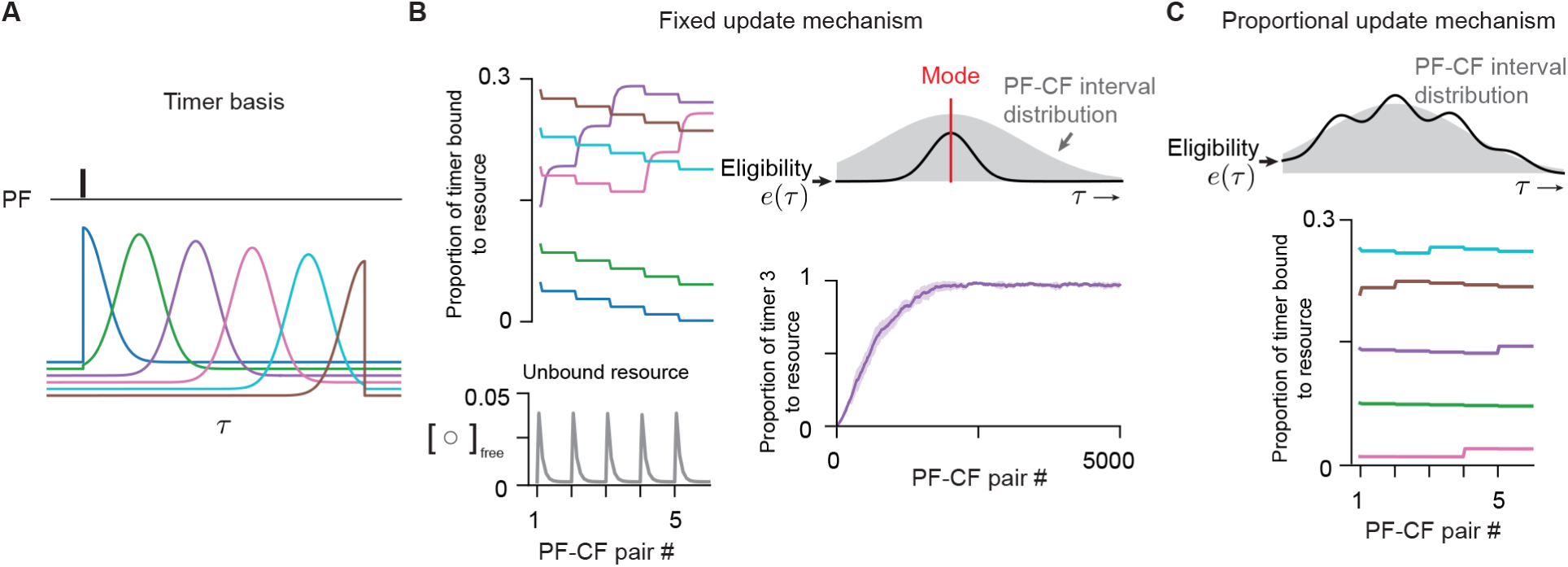
Further illustration of biochemical implementations of timer selection mechanism for temporal metaplasticity. **(A)** Simple bank of six timers for illustrating biochemical implementations of the timer selection mechanism. Peak times were spread evenly across 200 ms after parallel fiber (PF) input. **(B)** Biochemical implementation of the winner-take-all timer selection mechanism with limited resource and zero-order kinetics. ***Left***, the changes in concentrations of activated timer molecules with bound resource C_i_*-U_i_* (***top***), and the free (unbound) limited resource (activator) molecule (***bottom***), similar to **Fig. 5G_2_**, but shown for 5 PF-CF spike pair presentations. Colors correspond to timer activations in panel **A**. ***Top right***, eligibility window (*black*) picks out the mode (*red*) of the PF-CF spike interval distribution (*grey*). ***Bottom right***, mean ± 1 standard deviation over 10 simulations of the concentration of activated and bound timer C_3_*-U_3_*, whose activation contains the mode. **(C)** Biochemical implementation of the timer selection mechanism with proportional updates from plentiful resource and first-order kinetics. ***Top***, eligibility window (*black*) approximates the shape of the PF-CF interval distribution (*grey*). ***Bottom***, changes in concentrations of activated timer molecules with bound resource as a result of 5 PF-CF spike pair presentations. Colors correspond to timer activations in panel **A**.

