## Supplementary Information is available for this paper. for "Experience Alters the Timing Rules Governing Synaptic Plasticity and Learning"

### 1 Supplemental Methods

##### 1.1 Timer adjustment mechanism extracts the median of the parallel fiber-climbing fiber interval distribution

Here we show analytically that an ideal version of the timer adjustment mechanism results in an eligibility window that is exactly centered at the median of the parallel fiber-climbing fiber interval distribution. To do this, we assume that eligibility has a time course $e(\tau;\tau_{\text{peak}})$, where as in the main text $\tau$ indexes time after a PF input, and we let the parameter $\tau_{\text{peak}}$ define the location of peak eligibility. We further assume that the distribution of parallel fiber-climbing fiber intervals is defined over the interval $\tau \in [0, T]$, with probability given by a density function

$$p(\tau) = \mathbf{P}(\tau_{\text{CF}} = \tau), \qquad \text{(Equation S1)}$$

and cumulative distribution function

$$F(\tau) = \int_0^\tau \mathrm{d}\tau \; p(\tau). \qquad \text{(Equation S2)}$$

Consider the following temporal metaplasticity rule for tuning:

$$\tau_{\text{peak}} \leftarrow \tau_{\text{peak}} + \beta \Delta\tau_{\text{peak}}(\tau_{\text{CF}}), \qquad \text{(Equation S3)}$$

where the update is defined by

$$\Delta\tau_{\text{peak}}(\tau_{\text{CF}}) = \text{sgn}(\tau_{\text{CF}} - \tau_{\text{peak}}), \qquad \text{(Equation S4)}$$

so that any climbing fiber input that arrives after the peak timing moves the peak later, and any climbing fiber input that arrives before the peak timing moves the peak earlier. In this case, equilibrium will be reached if the tuning update, Equation S4, is zero on average. This occurs when

$$\begin{aligned}\mathbf{E}[\Delta\tau_{\text{peak}}] &= \int_0^T \mathrm{d}\tau\ p(\tau)\,\text{sgn}(\tau - \tau_{\text{peak}}) \\ &= \int_{\tau_{\text{peak}}}^T \mathrm{d}\tau\ p(\tau) - \int_0^{\tau_{\text{peak}}} \mathrm{d}\tau\ p(\tau) = 0,\end{aligned}$$

that is, for $\tau_{\text{peak}}$ such that

$$F(T) - F(\tau_{\text{peak}}) = F(\tau_{\text{peak}}).$$

This is solved when

$$\tau_{\text{peak}} = F^{-1}(1/2),$$

i.e., when $\tau_{\text{peak}}$ is the median timing (Figure 5H).

#### 1.2 Biochemical implementation of cascade of sequentially active timers

Here, we describe the implementation of a bank of molecular timers via a cascade of sequentially active elements. A reaction diagram is shown in Figure S8A (top). Briefly, the model contains three types of molecular species: species A, which is maximally activated by a parallel fiber input and is continuously degraded; a set of species $\{\mathrm{C}_m\}$ that form the temporal basis for eligibility; and a set of species $\{\mathrm{D}_m\}$ that are used to produce a delayed activation between subsequent $\mathrm{C}_m$'s (Figure S8A, bottom). The total concentration of active and inactive molecules was conserved by all reactions. The dynamical evolution of the fractional concentrations of these species are mathematically described by

$$\begin{aligned}\frac{\mathrm{dA}}{\mathrm{d}\tau} &= -k_{\mathrm{A}}^{-}\mathrm{A} \\ \frac{\mathrm{dC}_1^*}{\mathrm{d}\tau} &= k_{\mathrm{C},1}^{+}\frac{\mathrm{A}^{20}}{K_{\mathrm{A}}^{20} + \mathrm{A}^{20}}(1 - C_1^*) - k_{\mathrm{C},1}^{-}C_1^* \\ \frac{\mathrm{dD}_m^*}{\mathrm{d}\tau} &= k_{\mathrm{D},m}^{+}(1 - \mathrm{D}_m^*)C_{m-1}^* - k_{\mathrm{D},m}^{-}\mathrm{D}_m^* \\ \frac{\mathrm{dC}_{m+1}^*}{\mathrm{d}\tau} &= k_{\mathrm{C},m+1}^{+}\frac{\mathrm{D}_m^{*\,20}}{K_m^{20} + \mathrm{D}_m^{*\,20}}(1 - C_{m+1}^*) - k_{\mathrm{C},m+1}^{-}C_{m+1}^*,\end{aligned} \qquad \text{(Equation S5)}$$

where in the bottom two equations, $m > 1$, and asterisks indicate the active form of each species $\mathrm{C}_m$ and $\mathrm{D}_m$. The cascade could be of arbitrary length $M$. Parameter values are listed in Table S2. We note that the Hill coefficients are comparable to those which could result from an ultrasensitive process [S1].

#### 1.3 Biochemical implementation of coupling strength updates in multiple timer selection mechanisms

We assume a fixed basis of activated timers $\{C_m^*\}$. We model each timer as having a binding site $U_m$, which can bind to a conserved resource S. The binding site can be in an active state $U_m^*$ or an inactive state $U_m$, and in either state can be bound to the resource (Figure S8B; S is illustrated by a circle in the diagram). We denote the bound state with a circle, such that $U_m^{*\circ}$ represents the activated and bound state. Note that the state of the binding site does not affect the activation of the timer, i.e., the transition between the inactivated state $C_m$ and the active state $C_m^*$.

Each timer controls eligibility for plasticity when it is in its active state and bound to resource, i.e., $C_m^* U_m^{*\circ}$ or $C_m^* U_m^{\circ}$. We assume that there is an effective separation of timescales between the dynamics of the timer and the binding of the resource, so that on the timescale of timer dynamics,

$$\begin{aligned} [U_m^{*\circ}] &= [C_m U_m^{*\circ}](\tau) + [C_m^* U_m^{*\circ}](\tau) \approx \text{const.}, \\ [U_m^{\circ}] &= [C_m U_m^{\circ}](\tau) + [C_m^* U_m^{\circ}](\tau) \approx \text{const.}, \end{aligned} \qquad \text{(Equation S6)}$$

i.e., the total concentrations of timers with either an active and bound binding site or an inactive and bound binding site at any time $\tau$ are constant. The relative fraction of timers that have an active and bound binding state is then $[U_m^{*\circ}]/[C_m]_{\text{total}}$, where $[C_m]_{\text{total}}$ is the total concentration of timers in all 8 possible states: timer active/inactive, binding site active/inactive, resource bound/unbound. We assume conservation of all elements, so $[C_m]_{\text{total}}$ is constant. The concentration of active timer in the states that contribute to eligibility is then the total concentration of active timers, multiplied by the fraction in each of those states:

$$\begin{aligned} [C_m^* U_m^{*\circ}](\tau) &= \frac{[U_m^{*\circ}]}{[C_m]_{\text{total}}} [C_m^*](\tau), \\ [C_m^* U_m^{\circ}](\tau) &= \frac{[U_m^{\circ}]}{[C_m]_{\text{total}}} [C_m^*](\tau). \end{aligned}$$

Then, the eligibility window can be written

$$\begin{aligned} e(\tau) &= \sum_{m=1}^{M} k_m ([C_m^* U_m^{*\circ}](\tau) + [C_m^* U_m^{\circ}](\tau)) \\ &= \sum_{m=1}^{M} k_m \frac{[U_m^{*\circ}] + [U_m^{\circ}]}{[C_m]_{\text{total}}} [C_m^*](\tau). \end{aligned} \qquad \text{(Equation S7)}$$

If we define

$$u_m = \frac{[U_m^{*\circ}] + [U_m^{\circ}]}{[C_m]_{\text{total}}}, \qquad \text{(Equation S8)}$$

and we set the rate constants $k_m = 1/[C_m]_{\text{total}}$ for all $m$, then this corresponds to the equation used in the non-biochemical description of Figure $4D_1$ (STAR Methods, Equation 8).

As mentioned in the Main Text, the two implementations of the timer selection mechanisms—i.e., the fixed update and proportional update rules—can be thought of as resulting from the same molecular interactions (Figure S8B) in two different parameter regimes. Below, we present the implementation for the fixed update rule first, then show how the proportional rule can be derived as a special case.

##### 1.3.1 Fixed update rule

We assume that the binding reactions occur on a longer timescale than the timer dynamics. Thus, because the active state of each timer only occurs briefly relative to the inactive state, when considering binding to the resource S below, we can approximate all timers as being in the inactive state $\mathrm{C}_m$. Biologically, this separation of timescales might occur if, for example, activation of the $\mathrm{U}_m$ elements corresponds to a molecular tag.

For the fixed update form of the rule, we assume that the conserved resource that is necessary for coupling to eligibility is relatively limited. Timers bind to the resource S whenever their binding site is in the active state $\mathrm{U}_m^*$,

$$\mathrm{C}_m\mathrm{U}_m^* + \mathrm{S} \rightarrow \mathrm{C}_m\mathrm{U}_m^{*\circ}, \qquad \text{(Equation S9)}$$

putting them into the active and bound state. We assume that this reaction is enzyme-catalyzed with Michaelis-Menten kinetics,

$$\frac{\mathrm{d}[\mathrm{U}_m^{*\circ}]}{\mathrm{d}\tau} = k_{\mathrm{U}}^{+}[\mathrm{S}]_{\mathrm{free}}\frac{[\mathrm{U}_m^*]}{K_{m,\mathrm{U}}^{+} + [\mathrm{U}_m^*]}, \qquad \text{(Equation S10)}$$

using the definition of $[\mathrm{U}_m^{*\circ}]$ and $[\mathrm{U}_m^{\circ}]$ in Equation S6 above and that, on the timescale of binding, all timers are approximated as inactive so that,

$$[\mathrm{C}_m^*\mathrm{U}_m^{*\circ}] \approx 0, \text{ and } [\mathrm{C}_m^*\mathrm{U}_m^*] \approx 0,$$

and recall that, as noted above in Equation S6, $[\mathrm{U}_m^{*\circ}]$ and $[\mathrm{U}_m^{\circ}]$ are only constant on the timescale of the timer dynamics and not on the timescale of binding.

On the other hand, timers unbind from the resource when their binding site is inactivated,

$$\mathrm{C}_m\mathrm{U}_m^{\circ} \rightarrow \mathrm{C}_m\mathrm{U}_m + \mathrm{S}.$$

The unbinding dynamics are also enzyme-catalyzed with Michaelis-Menten kinetics,

$$\frac{\mathrm{d}[\mathrm{U}_m^{\circ}]}{\mathrm{d}\tau} = -k_{\mathrm{U}}^{-}\frac{[\mathrm{U}_m^{\circ}]}{K_{m,\mathrm{U}}^{-} + [\mathrm{U}_m^{\circ}]}, \qquad \text{(Equation S11)}$$

where we define

$$[\mathrm{U}_m^{\circ}] = [\mathrm{C}_m\mathrm{U}_m^{\circ}] + [\mathrm{C}_m^*\mathrm{U}_m^{\circ}] \approx [\mathrm{C}_m\mathrm{U}_m^{\circ}].$$

Finally, we assume that S is a conserved resource,

$$\frac{\mathrm{d}[\mathrm{S}]_{\mathrm{free}}}{\mathrm{d}\tau} + \sum_{m=1}^{M} \left( \frac{\mathrm{d}[\mathrm{U}_m^{*\circ}]}{\mathrm{d}\tau} + \frac{\mathrm{d}[\mathrm{U}_m^{\circ}]}{\mathrm{d}\tau} \right) = 0. \quad \text{(Equation S12)}$$

We model the total concentration of timers as large relative to the parameters $K^+_{m,\mathrm{U}}$ and $K^-_{m,\mathrm{U}}$ (the half-rate concentrations), so the reactions are rate-limited—i.e., in a saturated, zero-order regime.

The dynamics of temporal metaplasticity with this rule are as follows:

1. When a climbing fiber input arrives, timers that are active but have an inactive and unbound binding site have their binding sites activated,

   $$\mathrm{C}_n^*\mathrm{U}_n \overset{\mathrm{CF}}{\rightleftharpoons} \mathrm{C}_n^*\mathrm{U}_n^*,$$

   where we use the index $n$ to indicate the active timer at the time of the CF spike. We conceptualize this forward transition as a fast, potentially voltage-activated conformational change that we model as instantaneous. This switches timer $n$ into a state in which it can bind free resource, Equation S9. We treat the reverse reaction as relatively slow on the timescale of the binding dynamics.

2. Climbing fiber input also switches timers that are inactive and whose binding site is active and bound into a state with an inactive binding site,

   $$\mathrm{C}_m\mathrm{U}_m^{*\circ} \overset{\mathrm{CF}}{\rightleftharpoons} \mathrm{C}_m\mathrm{U}_m^{\circ} \quad \text{for all } m \neq n.$$

   For the fixed update rule, we model the forward reaction as instantaneous at the time of CF spike and the reverse reaction as also instantaneous a short time later (on the timescale of the timer dynamics). After the forward reaction, the timers will have their resource unbound according to Equation S11. As a result of the saturated zero-order kinetics, this short inactivation period of the binding site causes all the inactive timers to release approximately the same amount of resource, independent of the fraction that are bound,

   $$\frac{\mathrm{d}[\mathrm{U}^{\circ}_{m\neq n}]}{\mathrm{d}\tau} \approx -k^-_{\mathrm{U}}. \quad \text{(Equation S13)}$$

   (We note that as $[\mathrm{U}_m^{\circ}]$ becomes very small, the dynamics do become first-order.)

3. After it has become inactive, the timer $n$ is still in a state to take up resource. We assume that the enzyme catalyzing the binding reaction is also rate-limited, and becomes

   $$\frac{\mathrm{d}[\mathrm{U}_n^{*\circ}]}{\mathrm{d}\tau} \approx k^+_{\mathrm{U}}[\mathrm{S}]_{\mathrm{free}}. \quad \text{(Equation S14)}$$

   Thus, the timer $n$ binds the free resource that was been released by the inactive timers in the previous step.

4. From Equation S8, the above changes in $[\mathrm{U}_n^{*\circ}]$ and $[\mathrm{U}_{m\neq n}^{\circ}]$ correspond to changes in the coupling strengths $u_n$ and $u_{m\neq n}$ that mirror the idealized model in Main Text Equation 9 and Equation 10.

We simulated this biochemical implementation of the fixed update method for a set of 6 idealized molecular timers that tile 200 ms of time (Figure 5G$_2$, S9A,B). To generate the results on the right side of S9B, we used rate constants $k_\mathrm{U}^+ = 10$ and $k_\mathrm{U}^- = 0.2$, and let $K_{m,\mathrm{U}}^+ = K_{m,\mathrm{U}}^- = 0.001$. We simulated metaplasticity using parallel fiber-climbing fiber intervals drawn from a clipped normal distribution with mean $\mu = 80$ ms and standard deviation $\sigma = 50$ ms, normalized over the range $[0, 200]$ ms. For each PF-CF pair, we determined which timer was most activated at the time of the climbing fiber input and then simulated Equation S10, Equation S11 and Equation S12. We modeled the effect of the CF causing inactive timers to lose resource as lasting 50 ms, and simulated 0.95 s after the time of CF the spike before the next spike pair arrived. In the left panels of Figure S9B, we show the changes in total concentration of timer with resource bound, $[\mathrm{U}_m^{*\circ}]$, for the six timer activations in Figure S9A for 5 consecutive parallel fiber-climbing fiber pairs. One example pairing is shown in Figure 5G$_2$.

##### 1.3.2 Proportional update rule

The proportional update rule can be derived from the same molecular interactions as the fixed rule, under the following simplifying assumptions: (i) the resource is relatively abundant compared to the timers; (ii) the total concentration of timer is small enough that the unbinding of resource from timers with inactivated binding sites ($\mathrm{U}_m^{\circ}$) is not rate-limited (i.e., first order). We further assume the CF-triggered transition from $\mathrm{U}_m$ to $\mathrm{U}_m^*$ is reversed at the same rate as $\mathrm{U}_m^{\circ} \to \mathrm{U}_m^{*\circ}$.

The dynamics of temporal metaplasticity for the proportional rule are triggered by CF input similar to the fixed rule: CF input quickly activates the binding site of active timers, $\mathrm{C}_m^*\mathrm{U}_m \to \mathrm{C}_m^*\mathrm{U}_m^*$, and inactivates the binding site of inactive and bound timers, $\mathrm{C}_m\mathrm{U}_m^{*\circ} \to \mathrm{C}_m\mathrm{U}_m^{\circ}$. For the proportional rule, we model the reverse of these transitions as both occurring after a short time on the timescale of binding, whereas for the fixed rule, the transition for inactivated binding sites was much slower than the transition for activated binding sites.

With these assumptions, the binding reaction in Equation S10 for the timer $n$ that was active at the time of the CF spike becomes

$$\frac{\mathrm{d}[\mathrm{U}_n^{*\circ}]}{\mathrm{d}\tau} \approx k_\mathrm{U}^+[\mathrm{S}]_\mathrm{total}\frac{[\mathrm{U}_n^*]}{K_{m,\mathrm{U}}^+} = k_\mathrm{U}^{+\prime}([\mathrm{C}_n]_\mathrm{total} - [\mathrm{U}_n^{*\circ}]), \qquad \text{(Equation S15)}$$

where for the right hand equality we used that

$$[\mathrm{C}_n]_\mathrm{total} = [\mathrm{U}_n^{*\circ}] + [\mathrm{U}_n^*] + [\mathrm{U}_n] + [\mathrm{U}_n^{\circ}] \approx [\mathrm{U}_n^{*\circ}] + [\mathrm{U}_n^*],$$

where the approximation indicates that the CF spike activates approximately all of the binding sites $\mathrm{U}_n$ of the active timer at the time of the CF spike and inactivates approximately all of the bound binding sites $U^{*\circ}_{m\neq n}$ of the inactive timers, and where, because S is plentiful, we treat its concentration as effectively constant, and denote this concentration as $[\mathrm{S}]_{\mathrm{total}}$. Similarly, the unbinding reactions in Equation S11 for the inactive timers $C_m$ become

$$\frac{\mathrm{d}[\mathrm{U}^{\circ}_{m}]}{\mathrm{d}\tau} \approx -k^{-}_{\mathrm{U}}\frac{[\mathrm{U}^{\circ}_{m}]}{K^{-}_{m,\mathrm{U}}} = -k^{-\prime}_{\mathrm{U}}([\mathrm{U}^{\circ}_{m}]) \quad m \neq n. \qquad \text{(Equation S16)}$$

Using the definition of $u_m$ in Equation S8, Equation S15 and Equation S16 simplify to Main Text Equation 12. If the reaction rates are equal, $k^{+}_{\mathrm{U}} = k^{-}_{\mathrm{U}}$, then the sum of the coupling strengths is conserved, as assumed in the Main Text.

We simulated this biochemical implementation of the proportional update method using Equation S15 and Equation S16 similarly to the fixed updates (see above), with rate constants $k^{+\prime}_{\mathrm{U}} = k^{-\prime}_{\mathrm{U}} = 1$. Results for the same clipped normal distribution of parallel fiber-climbing fiber spike intervals are shown in Figure S9C (top). In the bottom panel of Figure S9C, we show the changes in concentration of the six eligibility coupling cofactors corresponding to the timer activations in panel A over a small number of parallel fiber-climbing fiber pair presentations.

##### 1.4 Fixed update method extracts the mode of the parallel fiber-climbing fiber interval distribution

We show here that, in an idealized setting, the timer selection mechanism with fixed updates, Equation 9 and Equation 10 in the main text, results in an eligibility window that consists only of the timer that is most frequently activated by the parallel fiber-climbing fiber interval distribution $p(\tau)$. Suppose that the timer bank has $M$ elements, and that $p_m$ defines the probability that element $m$ will be maximally activated (compared to the other elements) at the time of the climbing fiber spike, i.e.,

$$p_m = \int_0^T \mathrm{d}\tau\ \Theta(\mathrm{X}_m(\tau) > \mathrm{X}_k(\tau)\ \forall\ k \neq m)p(\tau), \qquad \text{(Equation S17)}$$

where $\Theta(\cdot)$ is the Heaviside function and $T$ is the maximum PF-CF spike interval that can be detected by the metaplasticity mechanism. Then, we will show that the eligibility window will tend toward a coupling weight of 1 for the timer $m^*$ for which $p_{m^*} > p_m$ for all $m$, and a coupling weight of 0 for all other timers.

Assuming that all timers have nonzero coupling weights, and ignoring the positivity constraint when $0 < u_m < \delta$ in Equation 10 of the main text, the expected change in a coupling weight due to a parallel fiber-climbing fiber pairing is

$$\begin{aligned}\mathbb{E}[\Delta u_m] &= (M-1)\delta p_m - \delta(1-p_m) \\ &= \delta(Mp_m - 1),\end{aligned} \qquad \text{(Equation S18)}$$

which is negative if $p_m < 1/M$, i.e., if $p_m$ is less than the probability of a pairing assuming a uniform distribution. If, as we assumed, the distribution has a unique mode, it is not uniform, and there will be at least one timer that has probability of activation $< 1/M$. Since for these timers, the average change in coupling weight will be negative, eventually their coupling weights will go to zero on average. We will say that all remaining timers with nonzero coupling weights have indices in the set $M_+$.

Then, for this smaller set of timers, the expected change in coupling weight due to a parallel fiber becomes

$$\begin{aligned}\mathbb{E}[\Delta u_m \mid m \in M_+] &= (|M_+| - 1)\delta p_m - \delta \left( \sum_{i \in M_+} p_i - p_m \right) \\ &= \delta \left( |M_+| p_m - \sum_{i \in M_+} p_i \right),\end{aligned} \quad \text{(Equation S19)}$$

since the coupling weight only decreases when climbing fibers occur during the activation of a timer with nonzero coupling weight. The expected weight change will be negative if

$$p_m < \frac{\sum_{m \in M_+} p_m}{|M_+|}, \quad \text{(Equation S20)}$$

where the right hand side is the average value of $p_m$ for timers in $M_+$. As before, there will be at least one timer with probability of activation that meets this criterion. All such timers will see their coupling weights eventually go to zero on average, reducing the set $M_+$. Eventually, there will be only two timers with nonzero weights, and the right hand side of Equation S20 will be the average of the two probabilities of activation for the two timers. Thus, the timer with the smaller probability of activation will see its coupling weight go to zero, and the timer with the larger probability of activation (and the largest overall) will see its coupling weight go to 1.

For an idealized basis consisting of a bank of timers each exclusively active for a short period of time tiling the time interval $[0, T]$, the time for which each timer is activated can be used to build a histogram from the probability distribution. The eligibility window resulting from the fixed update method will pick out the mode of the distribution when discretized in this way.

##### 1.5 Proportional update method causes eligibility to approximate the parallel fiber-climbing fiber interval distribution

For the timer selection mechanism with proportional updates according to Equation 12 in the main text, we can find the equilibrium eligibility coupling values, as for the timer adjustment mechanism above, by finding when the changes to the couplings are on average 0. As in the previous section, we assume that $p_m$, Equation S17, is the total probability that a climbing fiber spike will occur while timer $m$ is the most active of the set. Then, the average change to the coupling over parallel fiber-climbing fiber intervals is

$$\mathbb{E}[\Delta u_m] = \gamma \left( p_m(1 - \mathbb{E}[u_m]) - (1 - p_m)\mathbb{E}[u_m] \right),$$

which equals 0 when

$$\mathbb{E}[u_m] = p_m. \qquad \text{(Equation S21)}$$

For an idealized basis consisting of a bank of timers each exclusively active for a short period of time tiling the time interval $[0, T]$, the eligibility will effectively be a histogram of the probability distribution with bins given by the widths of the timer activations, which will approximate the probability density increasingly well as the timer activation widths get smaller.

#### 1.6 Cerebellar circuit model

Many circuit models of cerebellar plasticity exist, including some that capture many detailed features of data sets at a high level of detail [S2, S3]. Here, we chose a simpler, spiking model that enabled us to highlight key dynamical features of the interactions between temporal metaplasticity, parallel fiber-Purkinje cell synaptic plasticity, and eye movement behavior. To better understand the dynamics of temporal metaplasticity, we built a simple model of the circuit underlying oculomotor learning (Figure 4E). We modeled a representative Purkinje cell, which received spiking inputs from a set of $N_{\text{PF}} = 120$ parallel fibers, as well as a molecular layer interneuron (MLI). On each trial $k$ (1 s period), a spike train $\text{PF}_j^{(k)}$ was drawn independently for each parallel fiber $j$, according to a mean firing rate $\overline{\text{PF}}_j(t)$ in each time bin. The activity of the Purkinje cell was given by a weighted sum of the parallel fiber spike trains convolved with an exponential synaptic filter,

$$\text{PC}^{(k)}(t) = \sum_{j=1}^{N_{\text{PF}}} w_j^{(k)}(\text{PF}_j^{(k)}(t) * e^{-t/\tau_s}) - w_{\text{MLI}}\text{MLI}(t), \qquad \text{(Equation S22)}$$

where $w_j^{(k)}$ is the PF-PC weight on trial $k$, $\tau_s = 10\,\text{ms}$ is the synaptic time constant, and molecular layer interneuron activity was calculated as pooling PF inputs,

$$\text{MLI}(t) = \sum_{j=1}^{N_{\text{PF}}} \overline{\text{PF}}_j(t) * e^{-t/\tau_s}. \qquad \text{(Equation S23)}$$

**Associative plasticity model** The trial-to-trial change in weight at each parallel fiber-Purkinje cell synapse due to climbing fiber-triggered associative LTD was computed as

$$w_j^{(k+1)} = w_j^{(k)} + \Delta w_j^{(k)} - k(w_0 - w_j^{(k)}), \qquad \text{(Equation S24)}$$

which includes a decay to the baseline weight of $w_0$, and

$$\Delta w_j^{(k)} = -k_{\text{LTD}} \int \text{d}t\ (\text{PF}_j^{(k)}(t) * e(t)) \cdot \text{CF}^{(k)}(t) + k_{\text{LTP}} \int \text{d}t\ \text{PF}_j^{(k)}(t), \qquad \text{(Equation S25)}$$

where the integral is taken over the length of the trial, $\text{CF}^{(k)}(t)$ is a delta function at the location of a climbing fiber spike chosen from a distribution $r_{\text{CF}}$ (described in the sections below), and $e(t)$ is the eligibility window for plasticity (Figure 4B). The first term represents the associative CF-driven LTD, and the second represents nonassociative LTP.

We modeled eligibility for plasticity as governed either by the winner-take-all multiple timer selection temporal metaplasticity mechanism or the narrow window, single adjustable timer temporal metaplasticity mechanism. Concretely, for the winner-take-all mechanism, eligibility was defined as

$$e(t) = \sum_{m=1}^{M} u_m c_m(t), \qquad \text{(Equation S26)}$$

where $c_m(t)$ are the timer activations and $u_m$ are their coupling strengths to eligibility. We chose $M = 12$ timers with square wave activations that tiled 200 ms. Narrow window timer selection was simulated using a timer activation given by a Gaussian around the current peak eligibility time $\tau_{\text{peak}}$ with standard deviation 10 ms, and changes in $\tau_{\text{peak}}$ were given by Equation 6 in the Main Text with $\tau_l = \tau_r = 20\,\text{ms}$. We assume that metaplasticity operates over a much slower timescale than plasticity.

Climbing fibers were driven by contraversive (here, negative) retinal slip errors, delayed according to a distribution $d(\tau)$ with mean at 120 ms. During trial $k$, given a time course of retinal slip $\text{RS}^{(k)}(t)$, we generated a CF spike, $\text{CF}^{(k)}(t)$, by drawing a spike time from the distribution

$$\mathbf{P}(\text{CF}^{(k)} \text{ at time } t) \propto ([\text{RS}^{(k)}]_+ * d)(t), \qquad \text{(Equation S27)}$$

normalized so that climbing fibers on average fire at 1 Hz. For simplicity we picked one CF spike time per 1 s trial.

##### 1.6.1 Poisson firing model

We have shown that temporal metaplasticity depends on the statistics of the intervals between parallel fibers and subsequent climbing fibers. To generate an estimate for the type of distribution that would drive metaplasticity in a circuit like the one underlying oculomotor learning, we directly simulated the activity of a single Purkinje cell, based on spiking input from parallel fibers, and a lumped (i.e., averaged) population of $N_{\text{PC}} - 1$ Purkinje cells, whose activity was determined by an average firing rate, within the circuit model described above. We used the simple assumption that each parallel fiber $j$ fires as a homogeneous Poisson process with rate $\overline{\text{PF}}_j^{(k)}(t) \equiv \overline{\text{PF}}$. This models the variability one might expect to see in a parallel fiber on long timescales over many different kinds of behavior. We model the retinal slip signal as

$$\text{RS}^{(k)}(t) = -k_E \left( \frac{1}{N_{\text{PC}}} \text{PC}^{(k)}(t) + c\overline{\text{PF}}\xi(t) \right), \qquad \text{(Equation S28)}$$

where we used $k_E = 0.1\,(^\circ/\text{s})/(\text{sp/s})$ and $N_{\text{PC}} = 100$, $\xi(t)$ was a standard normal noise process, and $c = \sqrt{(1/N_{\text{PC}})(1 - 1/N_{\text{PC}})}$. The first term models the effect of the single model PC. The second term models the contribution to the eye movement output of the rest of the population of

$N_{\mathrm{PC}}$ Purkinje cells, modeled as a noise process with mean 0 and standard deviation $c$. In this way, there is always some small correlation of the PC to the error signal that drives learning. Plasticity was implemented as described above, and temporal metaplasticity was implemented with either the winner-take-all timer selection metaplasticity rule (Figure 4F) or the narrow timer adjustment rule.

We simulated the model over 10 blocks of 1,000,000 trials each for computational efficiency and to keep any small imbalances in the rates of LTD and LTD from potentially accumulating over the very long timescales of the simulations. We used a mean PF firing rate value of $\overline{\mathrm{PF}} = 0.477\,\mathrm{sp\,s^{-1}}$, corresponding to the same number of spikes per PF in 1 s as in the oculomotor learning simulation below. Weights were constrained to stay between a minimum value of 0 and and a maximum value of 5, and we initialized them at the start of each simulation block to a value of 2.5. $w_{\mathrm{MLI}}$ also set to 2.5 so that this value defined a baseline weight. We set the scale for plasticity by setting $k_{\mathrm{LTD}} = 0.03\,\mathrm{s^2/sp^2}$ and chose $k_{\mathrm{LTP}} = 3.3 \times 10^{-4}\,\mathrm{s\,sp^{-1}}$ empirically to minimize the buildup of weights, and included a small decay in the weights on every PF spike, with decay rate $1 \times 10^{-3}\,\mathrm{sp^{-1}}$. The rate of temporal metaplasticity was controlled for the multiple timer selection mechanism by a value of $\delta = 1 \times 10^{-4}$, which defined the size of the decrease in coupling strength for active timers. For the single timer adjustment mechanism, the rate of temporal metaplasticity was controlled by a maximum value of $\beta = 5 \times 10^{-4}$ (see Main Text, Equation 5), which defined how much peak eligibility moved for each PF-CF pairing. After each simulation block, we calculated the cumulative histogram of PF-CF intervals for each synapse, reset the weights to baseline, and stored the metaplasticity coupling strengths to carry over to the next block.

For both temporal metaplasticity mechanisms, the distributions of PF-CF intervals measured have a small peak around the mean of the true delay distribution, on top of a large uniform background (Figure 4I, bottom; S6A). By the end of the simulation, the eligibility window for plasticity had shifted to align with this peak, which occurred around 120 ms. This corresponded to a coupling strength of $u_7 = 1$ and $u_{m \neq 7} = 0$ at all synapses for the multiple timer selection model, and to a mean value of $\tau_{\mathrm{peak}} \approx 125\,\mathrm{ms}$ across the population for the single timer adjustment model. This is because, since firing is relatively uniform over time, when a climbing fiber spike occurs, it is contributing to a wide variety of timings across the population of parallel fibers.

##### 1.6.2 Oculomotor learning

To better compare to the experiments, we modeled training to increase the gain of the optokinetic reflex in the circuit model. We assume that an experimentally imposed external sinusoidal stimulus drives the parallel fiber population to fire in a structured manner. The external stimulus also strongly determines the retinal slip signal carried by the climbing fiber.

For this model, as above, we modeled both a representative Purkinje cell as well as a lumped population of Purkinje cells that contribute to the eye movement output. The population was modeled as

$$\overline{\mathrm{PC}}^{(k)}(t) = \sum_{j=1}^{N_{\mathrm{PF}}} \overline{w}_j^{(k)} \overline{\mathrm{PF}}_j(t) * e^{-t/\tau_s} - w_{\mathrm{MLI}}\mathrm{MLI}(t)$$
$$= \sum_{j=1}^{N_{\mathrm{PF}}} (\overline{w}_j^{(k)} - w_0)(\overline{\mathrm{PF}}_j(t) * e^{-t/\tau_s}), \quad \text{(Equation S29)}$$

where we set $w_{\mathrm{MLI}} = w_0$, and the parallel fiber firing rates $\overline{\mathrm{PF}}_j(t)$ are defined below. After each trial (i.e., sinusoidal cycle) $k$, we updated the population weights $\overline{w}_j$ according to the plasticity rule, Equation S24 to Equation S25. We modeled the single representative Purkinje cell as driven by parallel fiber spiking inputs $\mathrm{PF}_j^{(k)}(t)$ that were inhomogeneous Poisson processes, with rates given by $\overline{\mathrm{PF}}_j(t)$ and weights $w_j$. Then, we modeled the eye movement output as a combination of the population and the representative PC,

$$E^{(k)}(t) = k_E \left[ \left(1 - \frac{1}{N_{\mathrm{PC}}}\right) \overline{\mathrm{PC}}^{(k)}(t) + \frac{1}{N_{\mathrm{PC}}} \mathrm{PC}^{(k)}(t) \right] + g_0 S(t - 0.04), \quad \text{(Equation S30)}$$

where $S(t)$ is the optokinetic stimulus input, delayed by $\sim$40 ms as was characteristic of our experiments, and where $g_0 \approx 0.35$ is the baseline gain of the optokinetic reflex. Here we used $k_E = 5\,(^\circ/\mathrm{s})/(\mathrm{sp/s})$. The retinal slip error driving climbing fiber spikes was consequently

$$\mathrm{RS}^{(k)}(t) = S(t) - E^{(k)}(t). \quad \text{(Equation S31)}$$

The velocity of the optokinetic stimulus input $S(t)$ was a 1 Hz sinusoid with a peak velocity of $10\,^\circ\,\mathrm{s}^{-1}$, as in the behavioral experiments. Previous modeling work has suggested that granule cell firing forms a temporal basis set that would support a wide range of learned, time-varying responses [S4, S5, S6, S7, S8, S9, S10, S11, S2, S12, S13]. Consistent with this, cerebellar neurons have been shown to tile time during a trial of behavior [S14, S15], as also described for neurons in other brain areas [S16]. Motivated by this previous work, we modeled the response of the $j$th parallel fiber to the optokinetic stimulus as

$$\overline{\mathrm{PF}}_j(t) \propto [S(t - \Delta_j) - \theta]_+, \quad \text{(Equation S32)}$$

where $\theta = 9$ Hz and $[\cdot]_+$ represents positive rectification. The delays $\Delta_j$ were evenly spaced values between $-250$ ms and 750 ms. For simplicity, we allowed the activity of parallel fibers to wrap around in time. Thus, the parallel fiber responses formed a sequentially active bank of 120 rectified sinusoids that tiled the time of stimulus presentation (example spike raster in Figure 4H, left).

We simulated OKR adaptation for 1 h of simulation time using both an untuned plasticity rule representing the initial state before temporal metaplasticity (0 ms preference; Figure 4G, green) and a tuned (120 ms; Figure 4G, gold) plasticity rule representing the state after temporal metaplasticity, as found above. Weights were initialized to $w_0 = w_{\mathrm{MLI}} = 2.5$ and plasticity was simulated as described in the previous sections separately for the population weights $\overline{w}_j$ and the spiking PF weights $w_j$. Since temporal metaplasticity is much slower than plasticity, so we modeled the plasticity rule as not changing during the 1 h of OKR adaptation. For the untuned plasticity rule, we used learning rates $k_{\mathrm{LTD}} = 1.25 \times 10^{-2}\,\mathrm{s}^2/\mathrm{sp}^2$ and $k_{\mathrm{LTP}} = 2 \times 10^{-4}\,\mathrm{s\,sp}^{-1}$, and for the tuned plasticity rule we used $k_{\mathrm{LTD}} = 4 \times 10^{-2}\,\mathrm{s}^2/\mathrm{sp}^2$ and $k_{\mathrm{LTP}} = 6.4 \times 10^{-4}\,\mathrm{s\,sp}^{-1}$. We used the slightly smaller learning rates for the untuned plasticity rule to better model the behavior. The model reproduced the experimental finding of a delay in the time of the peak of the learned component of the eye movement response when using the untuned plasticity rule, relative to when using the tuned plasticity rule (Figure 4H; cf. Figure 2D).

###### 1.6.3 Cerebellar circuit model with a structured PF basis

To test whether the cerebellar circuit results depended upon having an uncorrelated PF basis state, or whether the same results would occur using a strongly correlated PF basis set, we built a model in which parallel fiber firing came from the same basis as in the oculomotor learning experiment. We modeled the retinal slip signal the same way as for the Poisson simulations, i.e., in Equation S28. Even with this structured basis, the PF-CF interval distributions show a clear peak at the time of the true physiological mean delay (Figure S6B, left, from a simulation using the winner-take-all temporal metaplasticity mechanism). Starting from an untuned plasticity rule, temporal metaplasticity in this model is able to tune the plasticity rule to select the true peak timing. Except for the choice of PF firing rates, the simulation was run in the same manner as for the Poisson basis.

**Table S1.** Reaction rates and threshold parameters for biochemical implementation of the accumulation-to-bound timer, related to STAR Methods

| Parameter | Value |
|---|---|
| $k^{+}_{A_1}$ | $10\,000\,s^{-1}$ |
| $k^{*}_{A_1 A_2}$ | 0.01 |
| $k^{-}_{A_1}$ | $2000\,s^{-1}$ |
| $k_{m,A_1}$ | 0.01 |
| $k^{-}_{A_2}$ | $100\,s^{-1}$ |
| $k_{A_2,\mathrm{off}}$ | $100\,s^{-1}$ |
| $k^{+}_{A_2}$ | $40\,s^{-1}$ |
| $k_{A_2 A^{*}_1}$ | 0.1 |
| $k^{+}_{B}$ | $5\,s^{-1}$ |
| $k_{BA^{*}_1}$ | 0.01 |
| $k^{-}_{B}$ | $7.5\,s^{-1}$ |
| $k_{B^{*}A_1}$ | 0.01 |
| $k^{+}_{C}$ | $10^{7}\,s^{-1}$ |
| $k_{CB^{*}}$ | 0.01 |
| $k^{-}_{C}$ | $2.5 \times 10^{7}\,s^{-1}$ |
| $k_{m,C}$ | 0.01 |

**Table S2.** Reaction rates and threshold parameters for biochemical implementation of the cascade of timers, related to STAR Methods

| Parameter | Value |
|---|---|
| $k_A^-$ | $100\,\mathrm{s}^{-1}$ |
| $k_{X,m}^+$ | $15\,000\,\mathrm{s}^{-1}$ |
| $k_{X,m}^-$ | $200\,\mathrm{s}^{-1}$ |
| $k_{D,1}^+$ | $100\,\mathrm{s}^{-1}$ |
| $k_{D,1}^-$ | $100\,\mathrm{s}^{-1}$ |
| $k_{D,m}^+$ for $m > 1$ | $80\,\mathrm{s}^{-1}$ |
| $k_{D,m}^-$ for $m > 1$ | $80\,\mathrm{s}^{-1}$ |
| $K_A$ | 0.6000 |
| $K_1$ | 0.3800 |
| $K_2$ | 0.3670 |
| $K_3$ | 0.3864 |
| $K_4$ | 0.3931 |
| $K_5$ | 0.3953 |
| $K_6$ | 0.3960 |
| $K_7$ | 0.3963 |
| $K_8$ | 0.3964 |
| $K_9$ | 0.3965 |
| $K_{10}$ | 0.3965 |
| $K_{11}$ | 0.3965 |
| $K_{12}$ | 0.3964 |
| $K_{13}$ | 0.3964 |
| $K_{14}$ | 0.3964 |
| $K_{15}$ | 0.3964 |
| $K_{16}$ | 0.3964 |
| $K_{17}$ | 0.3966 |
| $K_{18}$ | 0.3967 |
| $K_{19}$ | 0.3961 |
| $K_{20}$ | 0.3966 |
| $K_{21}$ | 0.3965 |
| $K_{22}$ | 0.3968 |
| $K_{23}$ | 0.3986 |
| $K_{24}$ | 0.4007 |
| $K_{25}$ | 0.4016 |
